# The genome sequence of the wild tomato *Solanum pimpinellifolium* provides insights into salinity tolerance

**DOI:** 10.1101/215517

**Authors:** Rozaimi Razali, Salim Bougouffa, Mitchell J. L. Morton, Damien J. Lightfoot, Intikhab Alam, Magbubah Essack, Stefan T. Arold, Allan Kamau, Sandra M. Schmöckel, Yveline Pailles, Mohammed Shahid, Craig T. Michell, Salim Al-Babili, Yung Shwen Ho, Mark Tester, Vladimir B. Bajic, Sónia Negrão

**Author notes:** shared first authors (Rozaimi Razali, Salim Bougouffa, Mitchell J. L. Morton, Damien J. Lightfoot). **Corresponding authors:** V.B. Bajic; King Abdullah University of Science and Technology (KAUST), Computational Bioscience Research Center (CBRC), Thuwal 23955-6900, Kingdom of Saudi Arabia; S. Negrão; King Abdullah University of Science and Technology (KAUST), Division of Biological and Environmental Sciences and Engineering, Thuwal, 23955-6900, Kingdom of Saudi Arabia; &966 8082569. **Author contribution** RR, SB, MJLM and DJL conceived and designed the analyses and managed particular components of the project. RR and SB performed the bioinformatics analyses, which included the compilation of genome scaffolds, annotation and genomic analyses. MJLM produced and analyzed the field data, performed the KO enrichment analysis and oversaw its biological context for data interpretation. RR performed the phylogenetic analyses. DJL produced the OrthoMCL results. SB and DJL performed the candidate gene analyses. IA and AK developed the computational tool Dragon Eukaryotic Analyses Platform (DEAP). ME and SA-B analyzed the inositol pathway and provided its biological context. STA performed the computational structure–function analysis of the I3PS protein. YP performed the *myo*-inositol quantification and analyzed the Na and K concentration, and SA-B provided its metabolic context. MS performed the field trial supervision and phenotypic data collection. MJLM, CTM and SMS prepared materials and undertook sequencing activities. DJL and YSH contributed to the bioinformatics and genomic analyses. RR, SB, MJLM, DJL and SN organized the manuscript, analyzed the data, and wrote the article. SN, MT and VBB designed the research, supervised the project and reviewed the article. All authors contributed to the writing of the paper. RR, SB, MJLM and DJL contributed equally.

## Abstract

*Solanum pimpinellifolium*, a wild relative of cultivated tomato, offers a wealth of breeding potential for several desirable traits such as tolerance to abiotic and biotic stresses. Here, we report the genome and annotation of *S. pimpinellifolium* ‘LA0480’. The ‘LA0480’ genome size (811 Mb) and the number of annotated genes (25,970) are within the range observed for other sequenced tomato species. We developed and utilized the Dragon Eukaryotic Analyses Platform (DEAP) to functionally annotate the ‘LA0480’ protein-coding genes. Additionally, we used DEAP to compare protein function between *S. pimpinellifolium* and cultivated tomato. Our data suggest enrichment in genes involved in biotic and abiotic stress responses. Moreover, we present phenotypic data from one field experiment that demonstrate a greater salinity tolerance for fruit-and yield-related traits in *S. pimpinellifolium* compared with cultivated tomato. To understand the genomic basis for these differences in *S. pimpinellifolium* and *S. lycopersicum*, we analyzed 15 genes that have previously been shown to mediate salinity tolerance in plants. We show that *S. pimpinellifolium* has a higher copy number of the inositol-3-phosphate synthase and phosphatase genes, which are both key enzymes in the production of inositol and its derivatives. Moreover, our analysis indicates that changes occurring in the inositol phosphate pathway may contribute to the observed higher salinity tolerance in ‘LA0480’. Altogether, our work provides essential resources to understand and unlock the genetic and breeding potential of *S. pimpinellifolium*, and to discover the genomic basis underlying its environmental robustness.

## INTRODUCTION

The *Solanum* section *Lycopersicon* is an economically important clade that consists of 14 species including the cultivated tomato *Solanum lycopersicum* (formerly *Lycopersicon esculentum*), which is the most economically important horticultural crop (Peralta *et al.*, 2005; Spooner *et al.*, 2005). This clade also includes *Solanum pimpinellifolium*, the closest wild relative of the cultivated tomato (The Tomato Genome Consortium, 2012; Aflitos *et al.*, 2014), that grows in the dry coastal regions of Peru, Ecuador, and northern Chile (Luckwill, 1943; Warnock, 1991; Peralta *et al.*, 2008). *S. pimpinellifolium* has a bushy growth type, small red fruits (~1.5 cm diameter) and is mostly facultative autogamous (Rick *et al.*, 1978). The dry coastal regions inhabited by this species are frequently exposed to brackish groundwater, salt-laden mist and other environmental conditions to which *S. pimpinellifolium* had to adapt, thus shaping its genetic and phenotypic variation (Rick *et al.*, 1977; Peralta and Spooner, 2000; Zuriaga *et al.*, 2009; Blanca *et al.*, 2012).

By virtue of its continued exposure to these challenging environmental conditions, *S. pimpinellifolium* exhibits a robustness that appears to have been lost in cultivated tomato during the domestication process (Miller and Tanksley, 1990; Tanksley and McCouch, 1997; Bai and Lindhout, 2007). Thus, *S. pimpinellifolium* is regarded as an important source of genes that can confer favorable stress-tolerance to cultivated tomato. For instance, breeding tomatoes with resistance to bacterial speck disease (caused by *Pseudomonas syringae*) was achieved though the introgression of the resistance (R) gene, *Pto,* from *S. pimpinellifolium* into commercial cultivars (Pitblado and Kerr, 1979; Pedley and Martin, 2003; Thapa *et al.*, 2015). Furthermore, horticultural traits of commercial tomato, such as fruit size, have been influenced by the introduction of *S. pimpinellifolium* alleles (as reviewed by Tanksley, 2004; Azzi *et al.*, 2015), some of which were identified by the molecular mapping of backcross populations developed from *S. pimpinellifolium* (Tanksley *et al.*, 1996). Additionally, numerous quantitative trait loci (QTLs) have been identified using *S. pimpinellifolium*, such as those for biotic stress (Salinas *et al.*, 2013; Chen *et al.*, 2014; Víquez-Zamora *et al.*, 2014; Ni *et al.*, 2017), abiotic stress (Villalta *et al.*, 2008; Lin *et al.*, 2010), fruit quality traits (Tanksley *et al.*, 1996; Chen *et al.*, 1999; Xiao *et al.*, 2008; Capel *et al.*, 2016), and other agronomic traits (Doganlar *et al.*, 2002; Cagas *et al.*, 2008; Nakano *et al.*, 2016). In fact, *S. pimpinellifolium* is known to have high salinity tolerance and is a promising source for improvement of salinity tolerance in cultivated tomato (Bolarín *et al.*, 1991; Villalta *et al.*, 2008; Estan *et al.*, 2009; Rao *et al.*, 2013; Rao *et al.*, 2015).

To drive research related to tomato and specifically increase the possibility of finding genes that confer favorable traits, the Tomato Genome Consortium published the high-quality genome sequence of *S. lycopersicum cv.* ‘Heinz 1706’, as well as a draft sequence of *S. pimpinellifolium* accession ‘LA1589’ (The Tomato Genome Consortium, 2012). The availability of the cultivated tomato genome has greatly contributed to several fields of research, such as the identification of candidate genes related to fruit development (Zhong *et al.*, 2013; Liu *et al.*, 2016), the development of SNP genotyping arrays (Sim *et al.*, 2012a; Sim *et al.*, 2012b; Víquez-Zamora *et al.*, 2014), the design of the CRISPR-cas9 gene-editing system (Brooks *et al.*, 2014), and the identification of loci associated with traits contributing to improved tomato flavor quality (Tieman *et al.*, 2017). Despite the availability of the draft genome sequence of *S. pimpinellifolium ‘*LA1589’, limited advances in the genetic studies of this species have been described so far (e.g. Kevei *et al.*, 2015), which may be attributed to the genome’s lack of contiguity (309,180 contigs), limited coverage, and lack of annotation. Thus, the availability of an improved genome sequence for *S. pimpinellifolium* is expected to overcome some of these limitations, and to provide opportunities for the discovery of new genes unique to wild germplasm within the *Lycopersicon* clade.

Here, we report a genome assembly of *S. pimpinellifolium* ‘LA0480’, an accession with high salinity tolerance, as shown in our field trial results. This genome assembly is a substantial improvement on the existing ‘LA1589’ assembly. The genome of *S. pimpinellifolium* ‘LA0480’ was sequenced using Illumina sequencing with approximately 197x coverage, and consists of 811 Mb with an N50 of 75,736 bp (Tables 1 and S4). Gene model annotation identified 25,134 protein-coding genes (Tables 1 and S9). Moreover, we release for the first time the Dragon Eukaryotic Analysis Platform (DEAP). The DEAP (pronounced DEEP) ‘Annotate’ module assigns annotation from multiple sources including the latest Kyoto Encyclopedia of Genes and Genomes (KEGG) database, UniProt and InterProScan. Additionally, the DEAP ‘Compare’ module provides a multitude of approaches to navigate, analyze and compare the different annotations, which enabled us to identify enrichments in genes involved in pathogen and stress responses. Furthermore, we demonstrate the utility of the *S. pimpinellifolium* genome sequence by exploring the genomic basis of its salinity tolerance. Our work will assist the discovery of new genes that can be introgressed into cultivated tomato to improve its salinity tolerance. We present field trial results that clearly show the higher salinity tolerance of *S. pimpinellifolium* compared to cultivated tomato, and we investigate the genomic basis for this superior performance under stress following a candidate gene approach. We find that genes encoding inositol-3-phosphate synthase (I3PS), a key enzyme involved in salinity response (Nelson *et al.*, 1998), have higher copy numbers in *S. pimpinellifolium* ‘LA0480’ compared with other, less salt tolerant, tomato species, suggesting that I3PS and the inositol pathway play an important role in salinity tolerance in ‘LA0480’. We also find that ABC proteins, a class of proteins with established roles in membrane transport, particularly of chloride and potassium, are enriched in *S. pimpinellifolium*, which might contribute to its higher salinity tolerance.

The improved genome of *S. pimpinellifolium* presented here will facilitate the use of this species in breeding programs to expand the genetic diversity of cultivated tomato. The *S. pimpinellifolium* ‘LA0480’ genome provides a highly valuable resource for comparative genomics, functional and evolutionary studies, and a background to dissect the genomic basis for traits unique to this wild species.

## RESULTS AND DISCUSSION

### *S. pimpinellifolium* ‘LA0480’ shows a higher salinity tolerance than cultivated tomato

To assess the salinity tolerance of *S. pimpinellifolium ‘*LA0480’ under field conditions, we phenotyped *S. lycopersicum* ‘Heinz 1706’ and *S. pimpinellifolium* ‘LA0480’ in a field trial under both control (non-saline) and saline conditions. From this field trial, we collected physiological measurements for both species specifically focusing on yield-related traits in the field that are the most relevant to breeders for downstream applications. We observed that the majority of traits were affected by salt stress in both genotypes (Table 1 and Figure 1), and that there were also clear differences in salinity tolerance (ST) index between genotypes across different traits. Strikingly, ‘LA0480’ ST values across all fruit-and yield-related traits were ~1.25 to 2.5 times greater than in ‘Heinz 1706’ (Table 1). The high ST for yield (total fruit fresh mass) in ‘LA0480’ relative to ‘Heinz 1706’ cannot be attributed to differences in fruit dimensions (fruit length and fruit diameter) or individual fruit mass (average fruit fresh mass), but appears to be the result of a marked increase in fruit number in response to salt in ‘LA0480’, whereas ‘Heinz 1706’ showed substantial reductions in all these traits under stress. That is, under salt stress, *S. pimpinellifolium* produced an increased quantity of fruit of similar size but reduced mass compared to control conditions; while *S. lycopersicum* produced fewer and smaller fruit under salt stress relative to control conditions. Interestingly, ST indices for shoot and total fresh and dry mass are 10-20% higher in ‘Heinz 1706’ than in ‘LA0480’. While this difference is modest, it is informative that this did not translate to enhanced yield maintenance under stress compared with control conditions. This observation highlights the importance of studying agronomically important traits directly, rather than relying solely on expedient proxies such as biomass measurements at the immature stage, which is in line with the findings of Rao *et al*. (2013). Higher ST for root traits in ‘LA0480’ than in ‘Heinz 1706’ provides an interesting correlation with high fruit-and yield-related ST, but further studies are required to understand this potential relationship.

Altogether, our field results confirm previous reports of high salinity tolerance in *S. pimpinellifolium* (Bolarín *et al.*, 1991; Villalta *et al.*, 2008; Rao *et al.*, 2013) and show, specifically, that ‘LA0480’ is more salt tolerant than ‘Heinz 1706’ in fruit and yield-related traits. These findings underline compelling physiological differences between the two accessions that merit further investigation and open possibilities to improve salinity tolerance in cultivated tomato. To establish the foundation for future research, we present the genome of *S. pimpinellifolium* accession ‘LA0480’ and investigate the genomic basis for its high salinity tolerance.

**Figure 1.**
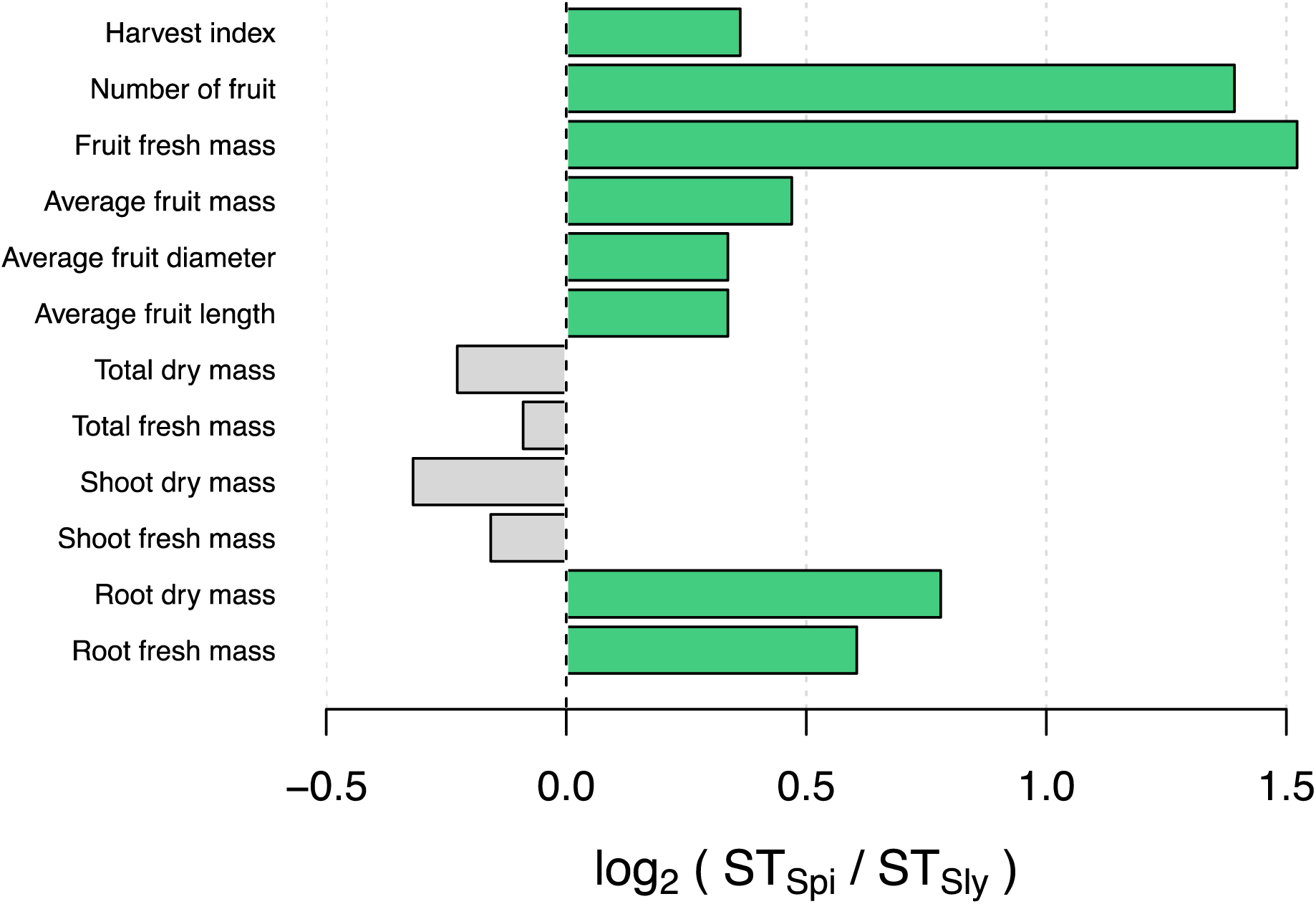
Comparison of *S. pimpinellifolium* and *S. lycopersicum* salinity tolerance (ST) indices across various traits measured in the field (log_2_ ratio) Traits for which the ST index is higher in *S. pimpinellifolium* and *S. lycopersicum* are in green and grey, respectively.

**Table 1.** Summary of field performance of *S. pimpinellifolium* and *S. lycopersicum* under control and saline conditions assessing various biomass and yield-related traits and their respective salinity tolerance (ST) index values for both species. ‘μ’ stands for mean and ‘se’ for standard error.

| Accession | Value | Root fresh mass (g) |  | Root dry mass (g) |  | Shoot fresh mass (g) |  | Shoot dry mass (g) |  | Total fresh mass (g) |  | Total dry mass (g) |  | Average fruit length (mm) |  | Average fruit diameter (mm) |  | Average fruit fresh mass (g) |  | Total fruit mass (g) |  | Yield (# of fruit) |  | Harvest Index |  |
| --- | --- | --- | --- | --- | --- | --- | --- | --- | --- | --- | --- | --- | --- | --- | --- | --- | --- | --- | --- | --- | --- | --- | --- | --- | --- |
|  |  | μ | ± se | μ | ± se | μ | ± se | μ | ± se | μ | ± se | μ | ± se | μ | ± se | μ | ± se | μ | ± se | μ | ± se | μ | ± se | μ | ± se |
| 'LA0480' | Control | 73.5 | 18.6 | 28.8 | 7.4 | 553 | 141 | 123 | 41.2 | 627 | 159 | 152 | 48.0 | 9.33 | 0.11 | 9.60 | 0.30 | 0.60 | 0.06 | 147 | 58.5 | 256 | 93.9 | 0.19 | 0.05 |
|  | Salt | 67.5 | 12.6 | 21.5 | 5.3 | 553 | 111 | 135 | 17.5 | 620 | 119 | 157 | 19.5 | 8.88 | 0.02 | 9.30 | 0.15 | 0.45 | 0.03 | 143 | 56.0 | 358 | 150 | 0.18 | 0.05 |
|  | ST | 0.92 |  | 0.75 |  | 1.00 |  | 1.10 |  | 0.99 |  | 1.03 |  | 0.95 |  | 0.97 |  | 0.75 |  | 0.97 |  | 1.40 |  | 0.95 |  |
| 'Heinz 1706' | Control | 19.3 | 1.3 | 7.8 | 1.3 | 142 | 20.4 | 36.9 | 7.7 | 162 | 20.8 | 44.6 | 8.3 | 41.9 | 3.1 | 27.8 | 1.58 | 26.8 | 1.44 | 439 | 77.9 | 19.5 | 3.63 | 0.73 | 0.05 |
|  | Salt | 11.6 | 3.9 | 3.4 | 1.5 | 159 | 86.1 | 50.5 | 26.2 | 170 | 89.7 | 53.9 | 26.7 | 31.6 | 3.7 | 21.3 | 3.21 | 14.5 | 1.25 | 149 | 40.1 | 10.4 | 2.95 | 0.55 | 0.12 |
|  | ST | 0.60 |  | 0.44 |  | 1.11 |  | 1.37 |  | 1.05 |  | 1.21 |  | 0.75 |  | 0.77 |  | 0.54 |  | 0.34 |  | 0.53 |  | 0.75 |  |
| ST<sub>Spi</sub>/ST<sub>Sly</sub> |  | 1.52 |  | 1.72 |  | 0.90 |  | 0.80 |  | 0.94 |  | 0.85 |  | 1.26 |  | 1.26 |  | 1.38 |  | 2.87 |  | 2.62 |  | 1.29 |  |
| log<sub>2</sub> (ST<sub>Spi</sub> /ST<sub>Sly</sub>) |  | 0.60 |  | 0.78 |  | -0.16 |  | -0.32 |  | -0.09 |  | -0.23 |  | 0.34 |  | 0.34 |  | 0.47 |  | 1.52 |  | 1.39 |  | 0.36 |  |

### Assembly and annotation of the *S. pimpinellifolium* reference accession ‘LA0480’ genome

The genome of *S. pimpinellifolium* ‘LA0480’ was sequenced using the Illumina HiSeq 2000 sequencing platform. We generated two paired-end libraries (insert sizes: 139 and 332 bp) and five mate-pair libraries (insert sizes: 2, 6, 8, 10 and > 10 kb) (Table S1), resulting in ~108 Gb and ~52 Gb of data, respectively, producing an estimated genome coverage of ~197x. The initial 160 Gb of raw data were processed to remove low quality sequences generating over 138 Gb of high quality data that were then assembled, scaffolded (Table S2) and gap-closed into 163,297 final scaffolds with an N50 of 75,736 bp and a total size of 811.3 Mb (Tables 2 and S4). The assembled genome size is within the expected range compared to closely related species such as *S. lycopersicum* (900 Mb) (The Tomato Genome Consortium, 2012) and *S. pennellii* (942 Mb - 1.2 Gb, (Bolger *et al.*, 2014a)). To assess the completeness of our genome assembly for all scaffolds above 1 kb, we used the Benchmarking Universal Single-Copy Orthologs database (BUSCO, (Simão *et al.*, 2015)). Of the 1,440 complete plant-specific single copy orthologs in the BUSCO database, we identified 1,375 (95.5%) orthologs in our assembly, denoting a high quality and nearly complete genome assembly (Table S5).

Analysis of the *S. pimpinellifolium* genome indicated that 59.5% of the assembled genome consisted of repetitive elements, with Long Terminal Repeats (LTR) retrotransposons of the Gypsy-type being the most abundant, comprising 37.7% of the assembled genome (Table S14; Figure S6). This result is consistent with the repeat content of the genomes of both *S. lycopersicum* (The Tomato Genome Consortium, 2012) and *S. pennellii* (Bolger *et al.*, 2014a). The *S. pimpinellifolium* assembly presented here represents a substantial improvement over the previously published *S. pimpinellifolium* draft genome, which contained 309,180 contigs and had an estimated genome size of 739 Mb (The Tomato Genome Consortium, 2012). We used a combination of *ab initio* prediction and transcript evidence supported by RNA-seq data from multiple tissues and conditions to annotate a total of 25,970 genes (25,134 protein-coding) (Table 2 and S9), with 21,016 genes (80.9%) assigned an AED score of less than, or equal to, 0.3, indicating that they are well supported. A BUSCO completeness score of 91.9% for the genome annotation (Table S8) was obtained, which is lower than the BUSCO results that we obtained for *S. lycopersicum* (99.0%) and *S. pennellii* (98.5%). This result is expected as the *S. lycopersicum* and *S. pennellii* genomes are more complete, as evidenced by their chromosome-level assemblies.

To investigate functional features of the protein-coding genes of *S. pimpinellifolium* and to compare with the protein-coding genes from closely related species (*S. lycopersicum*, *S. pennellii* and *S. tuberosum*), we developed DEAP, http://www.cbrc.kaust.edu.sa/deap/, which is an extension of Dragon Metagenomic Analyses Platform (DMAP, http://www.cbrc.kaust.edu.sa/dmap). The longest protein isoform of each gene was submitted to DEAP Annotate v1.0 for functional annotation (Table S10). Additionally, the longest protein isoform of each gene from the *S. lycopersicum* (NCBI annotation release 102, Nov 2016), *S. pennellii* (NCBI annotation release 100, Dec 2015) and *S. tuberosum* (NCBI annotation release 101, Jan 2016) genomes were annotated in the same manner.

**Table 2.** Genome assembly and annotation statistics for *S. pimpinellifolium* ‘LA0480’ in comparison to *S. pimpinellifolium* ‘LA1589’ (The Tomato Genome Consortium, 2012), *S. lycopersicum* (The Tomato Genome Consortium, 2012) and *S. pennellii* (Bolger *et al.*, 2014a).

| Species | Genome size (Mb) | Number of scaffolds | Longest scaffold (bp) | Scaffold N50 (bp) | Average scaffold length (bp) | Total number of predicted genes |
| --- | --- | --- | --- | --- | --- | --- |
| *S. pimpinellifolium* 'LA0480' | 811.3 | 163,297 | 893,636 | 75,736 | 4,968 | 25,970 |
| *S. pimpinellifolium* 'LA1589' * | 688.2 | 309,180 (contigs) | 80,806 (contigs) | 5,714 (contigs) | 2,226 | N/A |
| *S. lycopersicum* | 815.7 | 372 | 98,543,444 | 66,470,942 | 2,192,838 | 30,391 |
| *S. pennellii* | 926.4 | 12 | 109,333,515 | 77,991,103 | 77,202,205 | 32,519 |

\* *S. pimpinellifolium* 'LA1589' was assembled to contig level only

### Comparative genomics within the Solanaceae

To investigate the gene space of the *S. pimpinellifolium*genome, we undertook a comparative genomics approach to compare *S. pimpinellifolium* to three other related species: a second wild tomato (*S. pennellii*); cultivated tomato (*S. lycopersicum*); and the more distantly related cultivated potato (*S. tuberosum*) (Figure 2). OrthoMCL analysis revealed 14,126 clusters of orthologs (containing 78,973 proteins) that are common to all four species analyzed and may represent the core set of genes in *Solanum*. A total of 715 clusters (2,438 proteins) were identified as being specific to the three members of the *Lycopersicon* clade, while 4,028 proteins were determined to be specific to *S. pimpinellifolium*, including 682 protein-coding genes with paralogs (Figure 2) and 3,346 proteins with no identified homologs (Table S13).

Of particular interest is the identity of genes encoding the 644 proteins identified as being specific to the two wild tomato species, which are both described as being more tolerant to abiotic stresses than cultivated tomato (e.g. Bolger *et al.*, 2014a; Rao *et al.*, 2015). This increased tolerance may be due to retention of ancestral *Lycopersicon* genes that were lost during domestication of cultivated tomato. Within this set of wild tomato-specific genes, we identified 34 *S. pimpinellifolium* genes with high confidence functional annotations. Specifically, we identified genes with high homology to oxidoreductases (FQR1-like NAD(P)H dehydrogenase (SPi16852.1) and tropinone reductase I (SPi19065.1)), calcium sensors (calmodulin-like protein 3 (SPi15382.1) and WRKY transcription factors (SPi13765.1 and SPi20050.1) that may be involved in abiotic stress tolerance in ‘LA0480’. FQR1-like NAD(P)H dehydrogenases have been linked to salinity tolerance (Laskowski *et al.*, 2002; Song *et al.*, 2016), while tropinone reductase I has been suggested to play roles in salt stress and drought tolerance (Taji *et al.*, 2004; Shaar-Moshe *et al.*, 2015). The roles of WRKY transcription factors and calmodulins (reviewed by Chen *et al.*, 2012) in abiotic stress tolerance are not well defined, but numerous studies have suggested roles for these proteins in salt, drought, heat and cold tolerance (Reddy *et al.*, 2011; Chen *et al.*, 2012; Niu *et al.*, 2012; Virdi *et al.*, 2015).

**Figure 2.**
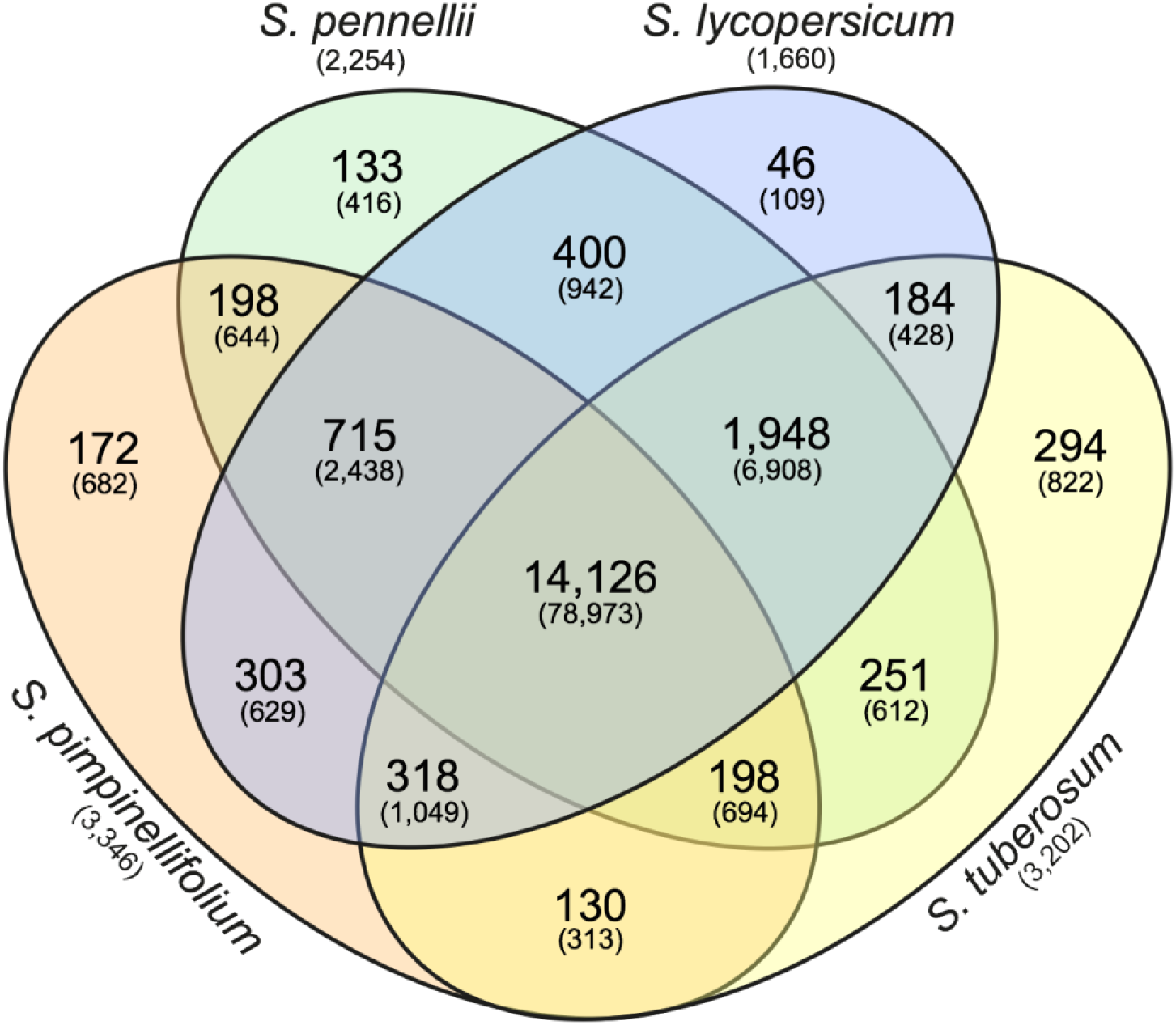
Identification of orthologous gene clusters in *S. pimpinellifolium, S. pennellii, S. lycopersicum* and *S. tuberosum*. The Venn diagram represents the number of protein-coding genes and gene clusters shared between, or distinct to, the indicated species. The number in each sector of the diagram indicates the number of homologous clusters and the numbers in parentheses indicate the total number of genes contained within the associated clusters. The numbers in parentheses below the species names indicate the number of species-specific singletons (genes with no homologs).

### Structural genomic variation between *S. pimpinellifolium* ‘LA0480’ and *S. lycopersicum* ‘Heinz 1706’

We investigated structural variation between the *S. pimpinellifolium* and *S. lycopersicum* genomes by identifying copy number variations (CNVs) due to duplication or deletion of genomic regions in either genome. We mapped *S. pimpinellifolium* and *S. lycopersicum* (SRA accession: SRR404081) short reads to the *S. lycopersicum* reference genome and identified regions of the *S. lycopersicum* genome with significantly increased coverage of either *S. pimpinellifolium* or *S. lycopersicum* reads after normalizing for differences in sequencing depth (Figure 3). CNV windows were identified as 276 bp sections of the *S. lycopersicum* genome where there was one log_2_-fold difference between the number of *S. pimpinellifolium* and *S. lycopersicum* mapped reads. CNV regions were called where there was at least 1,000 bp of contiguous CNV window coverage. We identified a total of 79,585 CNV regions, with 17,271 and 62,314 regions with higher and lower CNV, respectively, in *S. pimpinellifolium* (Table S15). The average length of these CNV regions is 3,024 bp, covering a total of 241 Mb (29.5%) of the *S. lycopersicum* genome. In *S. pimpinellifolium*, we observed substantially more low than high CNV regions, presumably because of the decreased mapping rate of the *S. pimpinellifolium* reads onto the *S. lycopersicum* reference genome as a result of sequence divergence between *S. pimpinellifolium* and *S. lycopersicum*. Thus, only regions with high CNVs in *S. pimpinellifolium* were analyzed further.

We identified 1,809 genes within these *S. lycopersicum* CNV regions, which is a comparable result to previous studies (Swanson-Wagner *et al.*, 2010; Cao *et al.*, 2011; Zheng *et al.*, 2011). Our analysis also indicated that 29.5% of the *S. lycopersicum* genome corresponds to CNV regions in *S. pimpinellifolium*. The proportion of the genome covered with CNV regions in this inter-species comparison is higher than what has been reported in intra-species comparisons (e.g. Belo *et al.*, 2010; Cao *et al.*, 2011; Yu *et al.*, 2011), where values are typically less than 5%. This difference is presumably due to the increased sequence divergence between the two tomato species investigated here.

As the identified CNVs represent regions of the genome that are substantially different between the two species, we investigated the *S. lycopersicum* genes that are within these CNV regions. We identified a total of 264 *S. lycopersicum* genes that may have a duplication of the corresponding regions in *S. pimpinellifolium* (Table S16), including genes that may play roles in abiotic or biotic stress tolerance. In particular, we identified one gene related to abiotic stresses tolerance such as drought and salt (LOC543714) (Islam and Wang, 2009); three genes related to leaf rust resistance (LOC101267807, LOC101268104 and LOC101254899) (Qin *et al.*, 2012); and two genes related to late blight resistance (LOC101264157 and LOC101258147) (Nowicki *et al.*, 2012). Additionally, we identified a number of *S. lycopersicum* transcription factors that may be duplicated in *S. pimpinellifolium* (e.g. LOC101259210, LOC101259230, LOC101262802 and LOC104649092). The identification of *S. lycopersicum* genes with roles in abiotic and biotic stress tolerance, that correspond to duplicated *S. pimpinellifolium* genes, provides candidates for further investigation as these genes that may underlie the increased stress tolerance in *S. pimpinellifolium*.

**Figure 3.**
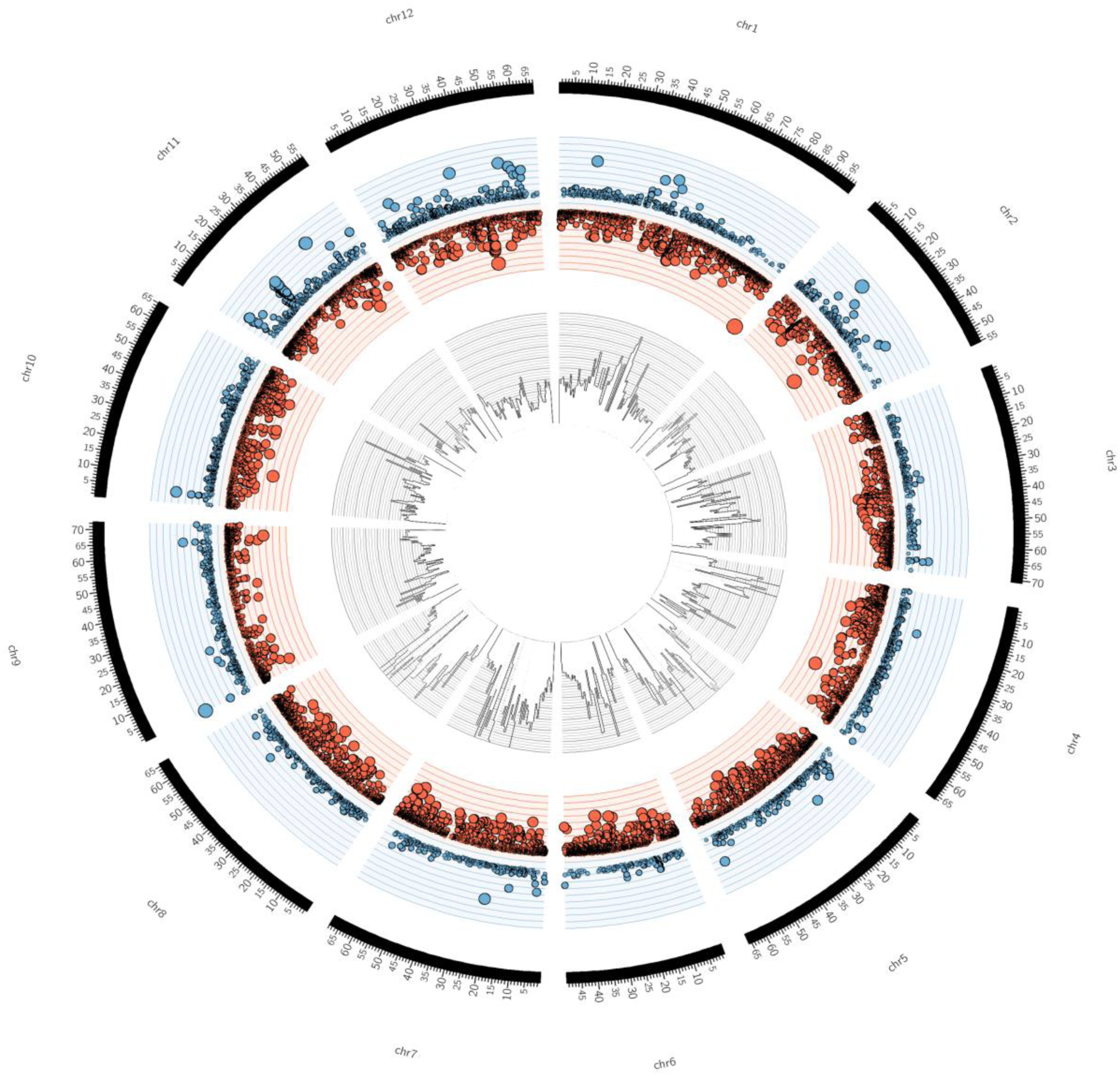
Circular representation of *S. pimpinellifolium* genome structure in comparison with *S. lycopersicum*. From the outside to the inside: The outer layer represents the 12 chromosomes of *S. lycopersicum,* with the axis scale in Mb. The second layer (blue/red) represents the scatter plot of copy number variant (CNVs) regions with blue and red circles denoting high and low copy variants, respectively in *S. pimpinellifolium* relative to *S. lycopersicum*. The size of the circles is proportional to the absolute value of the log_2_ CNV. The y-axis scale on the second layer corresponds to the log2 CNV ranging from -10 to 10. The innermost layer represents the histogram of SNPs in 1 Mb bins. The y-axis scale on the innermost layer represents the SNP distribution between the two species, which ranges from 0 to 19,095 SNPs.

### *S. pimpinellifolium* shows an enrichment in classes of genes related to stress responses

To identify classes of genes that are overrepresented in *S. pimpinellifolium*, with respect to *S. lycopersicum*, we annotated both genomes with KEGG Ontology (KO) terms using DEAP and performed an enrichment analysis (Figure 4). We discuss only those KO terms that are enriched in *S. pimpinellifolium* because the *S. lycopersicum* and *S. pimpinellifolium* genome assemblies have different levels of fragmentation, which could lead to the under-representation of some KO terms in the *S. pimpinellifolium* assembly. While correction for multiple testing using the Benjamini-Hochberg method detected only one significant enrichment, the KO analysis still highlights KO terms that are likely to be biologically relevant (on the basis of high fold change or absolute difference). Therefore, we discuss the top-ranking terms.

**Figure 4.**
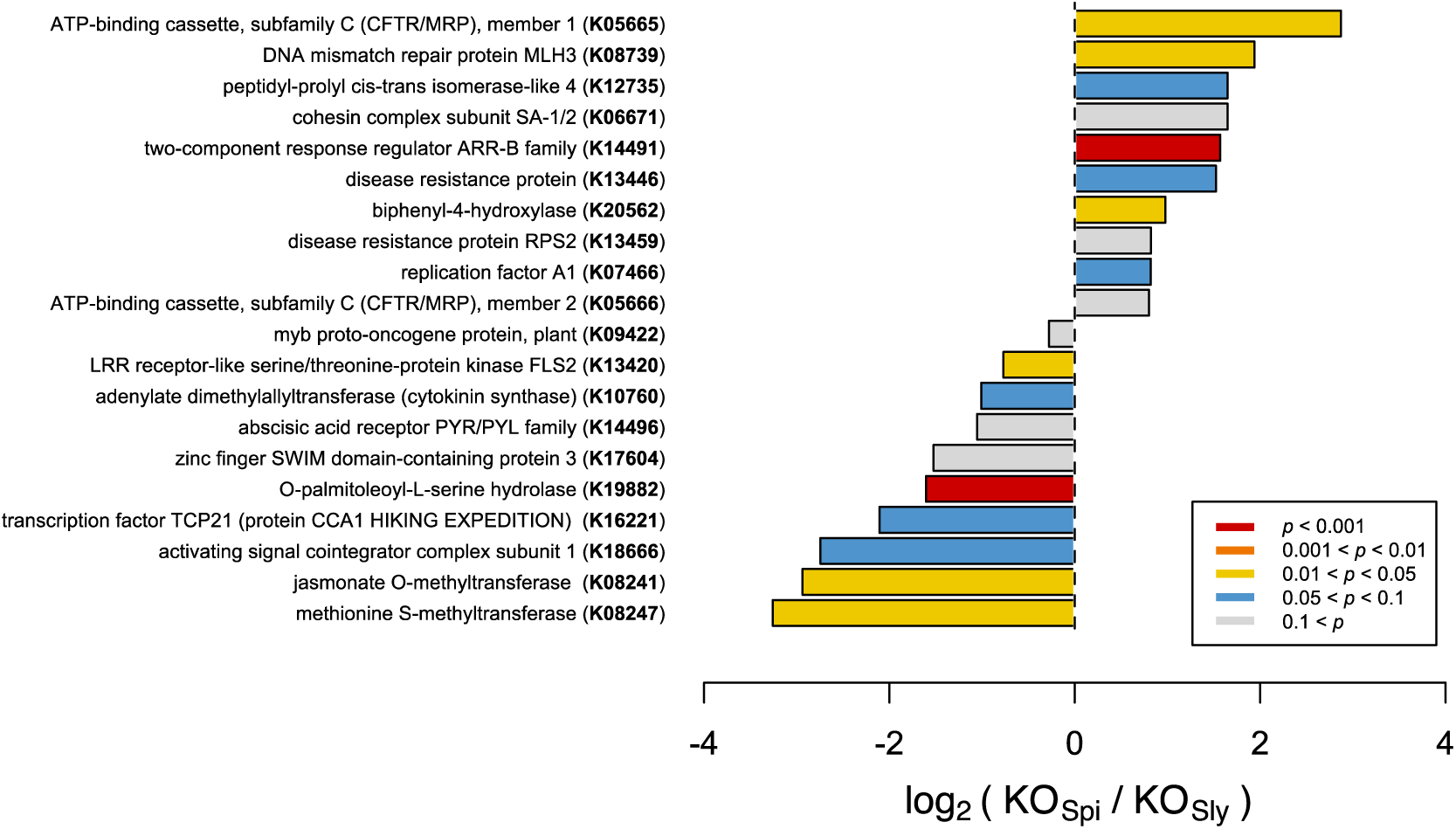
Comparison of KO term frequency in *S. pimpinellifolium* (KO_Spi_) and *S. lycopersicum* (KO_Sly_) genomes, presented as the ratio on a log_2_ scale. Bars are color-coded based on the *P* values from a Fisher’s exact test-based enrichment analysis (corrected for multiplicity using the Bonferroni method); the top 20 entries with the highest *P* values are presented. Entries are ordered based on log_2_ values.

Our analysis detected multiple KO terms that are significantly enriched in *S. pimpinellifolium* with respect to *S. lycopersicum*, several of which, according to KEGG classification, pertain to biological processes associated with biotic and abiotic stress tolerance, such as ‘two-component response regulator ARR-B family’ (K14491; *P* value < 3E-05), ‘biphenyl-4-hydroxylase’ (K20562; *P* value < 0.025), ‘DNA mismatch repair protein MLH3’ (K08739; *P* value < 0.035) and ‘ATP-binding cassette, subfamily C (CFTR/MRP), member 1’ (K05665, *P* value < 0.04). To elucidate the downstream biological relevance of such enrichments, we further investigated the functions of genes annotated with the corresponding KOs.

The KO term ‘two-component response regulator ARR-B family’ (K14491) denotes members of the Type-B response regulators (RR-B), a class of transcription factors that are the essential and final effectors in cytokinin (CK) signal transduction (Mason *et al.*, 2005). We observed 51 and 18 occurrences of this KO term in *S. pimpinellifolium* and *S. lycopersicum*, respectively. RR-Bs have been implicated in pathogen defense, acting as a bridge between cytokinin signaling and salicylic acid and jasmonic acid immune response pathways (Choi *et al.*, 2010; Argueso *et al.*, 2012). Moreover, cytokinins are involved in salinity responses (Tran *et al.*, 2007; Ghanem *et al.*, 2008; Mason *et al.*, 2010), with overexpression of cytokinin biosynthesis genes in *S. lycopersicum* shown to increase salinity tolerance (Ghanem *et al.*, 2011; Žižková *et al.*, 2015). These results suggest that the apparent expansion of RR-Bs in *S. pimpinellifolium* could contribute towards the increased pathogen resistance and stress tolerance of the species.

The ‘Biphenyl-4-hydroxylase’ (K20562) KO term was detected 32 and 17 times in the *S. pimpinellifolium* and *S. lycopersicum* genomes, respectively. Biphenyl-4-hydroxylases (B4H) have only recently been identified and cloned in rowan (*Sorbus aucuparia*) and apple (*Malus* spp.) and were characterized as cytochrome P450 736A proteins that catalyze the 4-hydroxylation of a biphenyl scaffold towards the biosynthesis of biphenyl phytoalexins such as aucuparin in response to pathogen attack (Khalil *et al.*, 2013; Sircar *et al.*, 2015). Research into biphenyl phytoalexins is somewhat scarce, possibly due to the absence of B4H in the model organism Arabidopsis, with most studies restricted to the Malinae subtribe of the subfamily Amygdaloideae (e.g. apple and pear) (Kokubun *et al.*, 1995; Hüttner *et al.*, 2010; Chizzali and Beerhues, 2012; Chizzali *et al.*, 2012; Sircar *et al.*, 2015; Chizzali *et al.*, 2016). As such, the observed presence and, indeed, expansion of B4H-related genes in *S. pimpinellifolium,* could be related to the increased pathogen resistance of *S. pimpinellifolium* and represents an interesting target for further studies.

In Arabidopsis, AtMLH3 (MutL protein homolog 3) regulates the rate of chromosome crossover during meiosis in reproductive tissues (Franklin *et al.*, 2006; Jackson *et al.*, 2006). We identified 11 and 3 occurrences of the corresponding KO term, ‘DNA mismatch repair protein MLH3’ (K08739), in the *S. pimpinellifolium* and *S. lycopersicum*, genomes respectively. A recent study on *Crucihimalaya himalaica*, an Arabidopsis relative that grows in the extreme environment of the Qinghai-Tibet Plateau, showed that the *C. himalaica* MLH3 homologue was under strong positive selection and may play a role in the repair of DNA damage caused by high UV radiation (Qiao *et al.*, 2016). This could point towards a role for MLH3-like genes repair of DNA damage (e.g. ROS-induced) caused by abiotic stress in *S. pimpinellifolium*.

‘ATP-binding cassette, subfamily C (CFTR/MRP), member 1’ (K05665) is enriched in *S. pimpinellifolium*, which has seven occurrences of this KO term against one in *S. lycopersicum*, suggesting an expansion of the ATP-binding cassette subfamily C (ABCC) protein superfamily in the wild tomato. ABC proteins encode transmembrane transporters and soluble proteins with crucial functions, and are ubiquitous across all kingdoms of life having a particularly high presence in plants (Andolfo *et al.*, 2015). ABCCs have been implicated in various transport processes in plants, such as vacuolar compartmentalization of glutathione conjugates, glucuronides and anthocyanins, as well as ATP-gated chloride transport, and the regulation of ion channels in guard cells (Martinoia *et al.*, 2002; Goodman *et al.*, 2004; Klein *et al.*, 2006; Suh *et al.*, 2007; Verrier *et al.*, 2008). Although the function of ABC proteins is difficult to determine from sequence similarity alone, we noted that the sole protein annotated with this KO in *S. lycopersicum*, namely XP_004248540, bears greatest sequence identity to Arabidopsis MRP9 (or ABCC9) and human SUR2 (sulfonylurea receptor 2), which are regulators of potassium channel activity (Rea, 2007). Because this class of genes has established roles in membrane transport, particularly of chloride and potassium, we hypothesize that the high number of ABCC annotated proteins in *S. pimpinellifolium* might contribute to its higher salinity tolerance. However, further analyses are required to determine the precise functions of these proteins and the extent of their involvement in such processes.

To complement the results of our comparative genomics analyses, we also undertook a literature-guided approach whereby genes with established roles in salinity tolerance were examined.

### Analysis of candidate genes that may confer salt tolerance

Given the higher salinity tolerance of *S. pimpinellifolium* and the broad and substantial knowledge of genes that contribute to salinity tolerance in tomato and other related species, we undertook a candidate gene (CG) approach to identify potentially important genes in *S. pimpinellifolium*. Based on the literature search summarized by Roy *et al.* (2014), we selected 15 CGs of primary interest (Table 3 and S17) that have been overexpressed in at least one plant species and were quantifiably shown to increase phenotypic performance under salt-stress conditions. Based on these CGs, we identified the corresponding *S. pimpinellifolium* orthologs based on published functional reports (e.g. *SlNNX1* (Gálvez *et al.*, 2012), OrthoMCL grouping (Figure 2), and reciprocal BLAST-P (Figure S9). Only CGs with orthologs that met these stringent criteria were considered for further analysis.

**Table 3.**
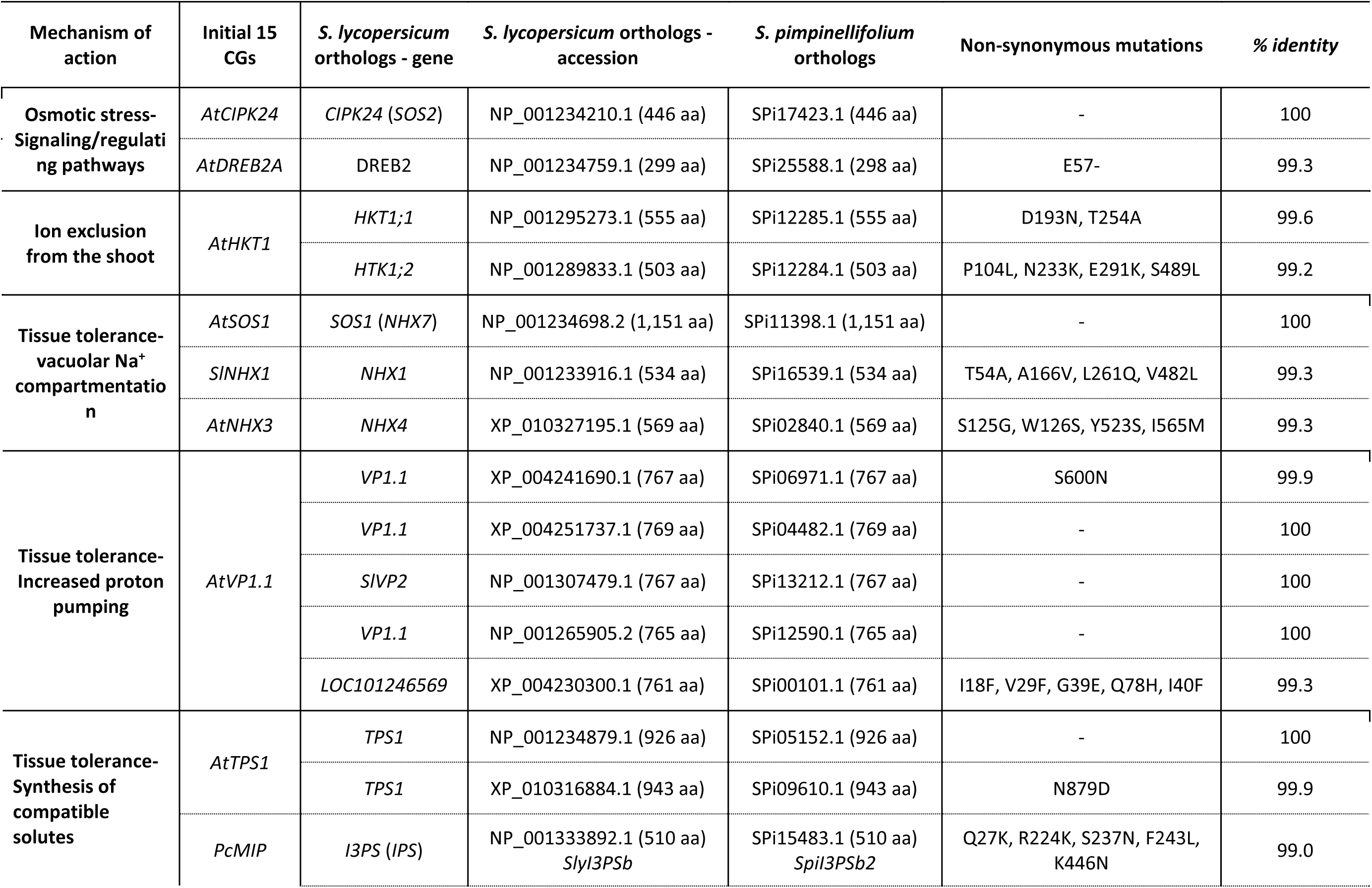
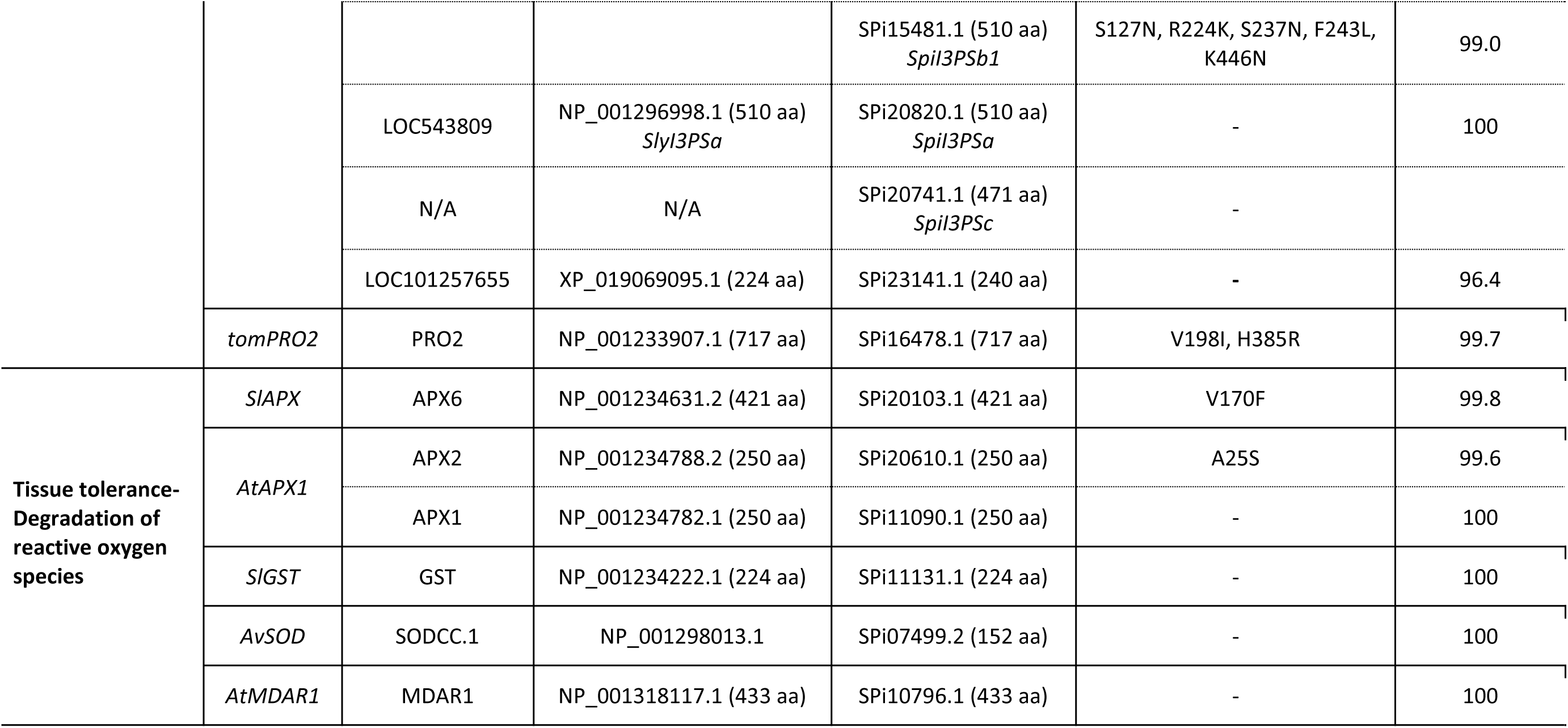
List of candidate genes for salinity tolerance in *S. pimpinellifolium* and *S. lycopersicum*. The source CG column lists the initial 15 candidate genes (for sequence accessions see Table S17). Gene symbol and full name were taken from the NCBI gene database. The non-synonymous mutations column and percentage identity columns are based on comparisons between the *S. lycopersicum* candidates and their respective homologs in *S. pimpinellifolium*.

We identified 24 putative orthologs in the *S. lycopersicum* genome that matched the selected 15 CGs. The *AtAVP1.1* gene has five orthologs in *S. lycopersicum, PcMIP* has three, while *AtHTK1, AtTPS1*, and *AtAPX1* have two orthologs each. The remaining ten CGs have one-to-one orthologous relationships. After establishing these *S. lycopersicum* orthologs, we investigated potential orthologs in *S. pimpinellifolium* (Table 3) and *S. pennellii* (Table S17; Figures S10-24). All of the *S. lycopersicum* genes have an identical number of orthologs in *S. pimpinellifolium* and *S. pennellii* except for the *inositol-3-phosphate synthase* (*I3PS*) gene. In terms of percentage identity, we observed a high similarity between the *S. lycopersicum* candidates and the corresponding orthologs in *S. pimpinellifolium* (> 99%, with 11 out of 24 reaching 100% similarity).

In the *S. lycopersicum* genome, we identified two copies of *I3PS* (*SlyI3PSa* and *SlyI3PSb*) as well as a truncated pseudogene (*LOC101257655*), while in the *S. pimpinellifolium* genome we identified four copies of *I3PS* as well as a truncated pseudogene. *S. pimpinellifolium* harbors one copy of *SpiI3PSa,* with 100% identity to *SlyI3PSa*, two copies of *SpiI3PSb* (*SpiI3PSb*1 and *SpiI3PSb*2), with more than 99% identity to *SlyI3PSb*, as well as *SpiI3PSc* (Table 3). At the DNA level, this fourth copy, *SpiI3PSc*, which is supported by RNA-seq evidence, is highly similar to the *SpiI3PSb* genes but contains two short frameshifts (not shown). As such, *SpiI3PSc* putatively encodes a protein product that is shorter than the other *SpiI3PS* genes due to the deletion of 36 amino acid residues.

To further investigate the relationships between the tomato *I3PS* genes, we built a DNA-based phylogenetic tree of the *I3PS* gene family using seven species of the Solanaceae, with Arabidopsis *MIPS* genes utilized as outgroups (Figures 5 and S25). Our results show a clear separation of the gene family into three distinct clades, namely the Arabidopsis, the “A” type and the “B” type clades, with the separation between the two gene types in Solanaceae being supported by high bootstrap values. In the “A-clade”, we observed a single-copy *I3PSa* gene for all Solanaceae species except for tobacco, which has two *IP3Sa* genes. The “B-clade” includes a single-copy of the *I3PSb* gene for all the Solanaceae species except for tobacco and *S. pimpinellifolium*, which both harbor two copies. This clade also contains *SpiI3PSc* (SPi20741), which appears to be a divergent *I3PSb* gene grouping with *S. pennellii*. However, the placement of these two sequences is unclear, as indicated by the low bootstrap value of 0.56. While the exact placement of *SpiI3PSc* and *SpeI3PSb* within the “B-clade” is unclear, alignment of the protein sequences (Figure S26) suggests that SpiI3PSc is an atypical form of the I3PS protein.

**Figure 5.**
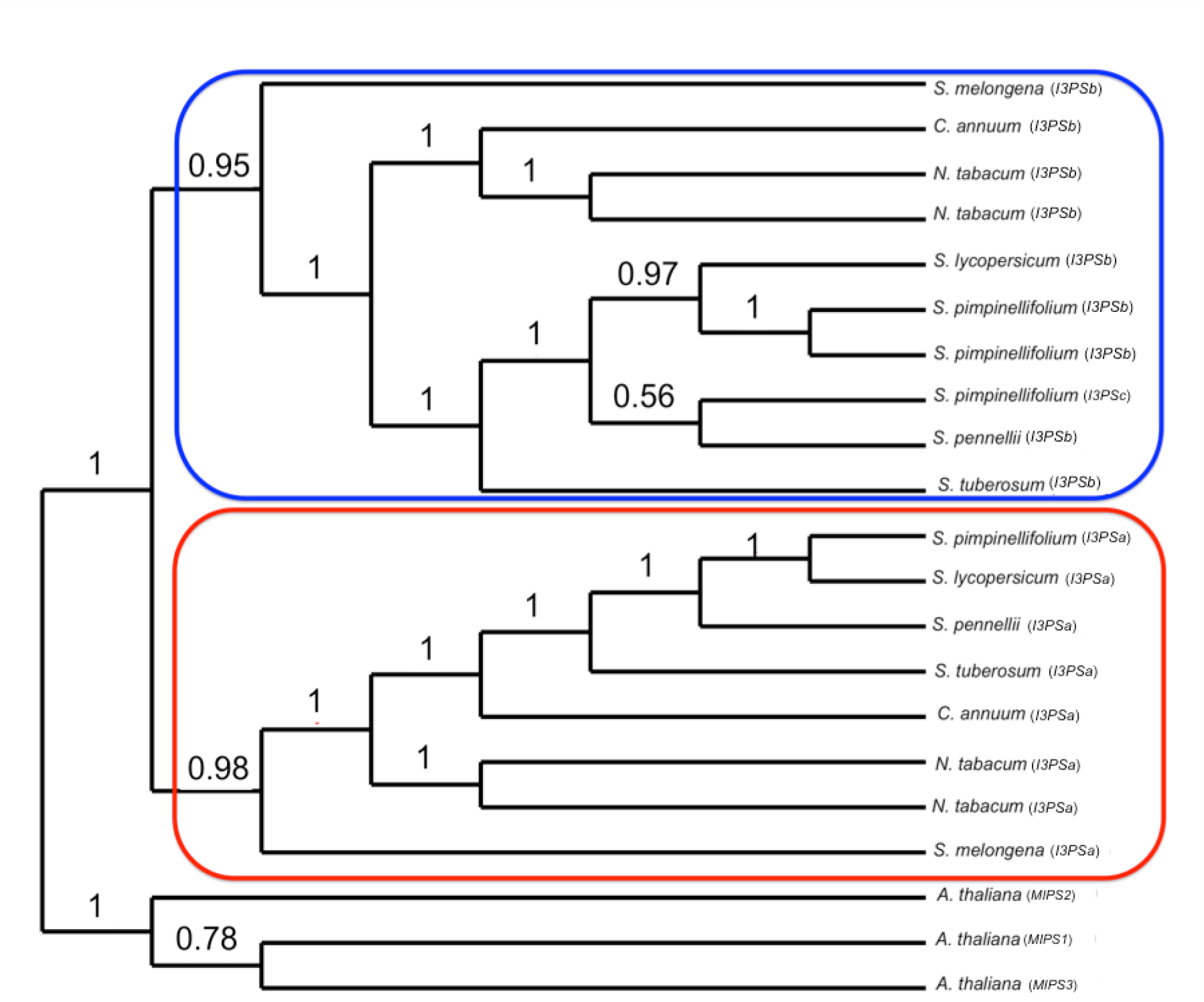
Phylogenetic analysis of the inositol-3-phosphate synthase (*I3PS*) gene family in the Solanaceae family. Node values represent the percentage of 100 bootstrap replicates that support the topology. The *I3PSa* and *I3PSb* genes are encircled in red and blue, respectively. *A. thaliana* MIPS genes were used as outgroups.

I3PS (EC:5.5.1.4) is a key enzyme in the inositol phosphate metabolism, which contributes to cell wall and membrane biogenesis, generates second messengers and signaling molecules, and provides compounds involved in abiotic stress response, phosphate storage in seeds, etc. (Bohnert *et al.*, 1995). I3PS is a NAD^+^-dependent enzyme that catalyzes the first step in the production of all inositol-containing compounds by converting glucose-6-phosphate (Glc6P) to D-*myo*-inositol-3-phosphate (Ins3*P*) (Majumder *et al.*, 2003; Stieglitz *et al.*, 2005), which is subsequently dephosphorylated by the inositol monophosphatase (EC:3.1.3.25) enzyme to *myo*-inositol (Stieglitz *et al.*, 2007). *Myo*-inositol is the substrate of the phosphatidylinositol synthase (PIS) (EC:2.7.8.11, CDP-diacylglycerol--inositol 3-phosphatidyltransferase) that forms the phospholipid phosphatidyl inositol (PtIns), an abundant phospholipid in non-photosynthetic membranes (Harwood, 1980; Boss and Im, 2012). The inositol moiety of PtIns can be targeted at the 3, 4 or 5 positions by specific kinases, leading to a variety of polyphosphoinositides, such as PtdIns3P, PtdIns4P, PtdIns5P, PtdIns(4, 5)*P*_*2*_, PtdIns(3, 5)*P*_2_ and PtdIns(3, 4)*P*_2_(Boss and Im, 2012). Phosphoinositides are involved in different cellular and developmental processes and contribute responses to various stresses. For instance, it was shown that the overexpression of phosphatidylinositol synthase in maize leads to increased drought tolerance by triggering ABA biosynthesis and modulation of the lipid composition of membranes (Liu *et al.*, 2013). *Myo*-inositol is also a precursor of soluble signaling molecules, such as Ins*P*_6_ (*myo*-Inositol hexakisphosphate also known as phytate) that acts as a second messenger triggering the release of Ca^2+^ from intracellular stores in guard cells (Lemtiri-Chlieh *et al.*, 2003), as well as ascorbic acid, a powerful reducing agent that is involved in scavenging reactive oxygen species under stress (reviewed by Akram *et al.*, 2017). Moreover, this compound plays a pivotal role in salinity tolerance by promoting the accumulation of its derivatives, such as D-pinitol and D-ononitol, as compatible solutes and thus protecting cells from osmotic imbalance (e.g. Nelson *et al.*, 1998; Nelson *et al.*, 1999). The accumulation of compatible solutes in the cell cytosol is critical for tissue tolerance, a key mechanism that involves the sequestration of Na^+^ ions in the vacuole (Tester and Davenport, 2003; Munns and Tester, 2008).

Introgression of the *I3PS* gene (*PcINO1*) from *Porteresia coarctata,* a halophytic wild rice, into rice, tobacco and Indian mustard has been reported to enhance the salinity tolerance of these species by substantially increasing inositol levels (Majee *et al.*, 2004; Das-Chatterjee *et al.*, 2006). Likewise, we suggest that the additional *SpiI3PS* gene copies identified in the wild tomato, *S. pimpinellifolium*, may contribute to its higher salinity tolerance when compared to cultivated tomato. However, further studies are necessary to validate the relative importance of each copy of *SpiI3PS* in *S. pimpinellifolium*.

### Assessing the role of I3PS in salinity tolerance of *S. pimpinellifolium*

To investigate if the four copies of *I3PS* in *S. pimpinellifolium* are functional, we first aligned the sequences of the eight I3PS proteins identified in the *Lycopersicon* species, namely two copies from *S. lycopersicum* (SlyI3PSa and SlyI3PSb), two copies from *S. pennellii* (SpeI3PSa and SpeI3PSb) and four copies from *S. pimpinellifolium* (SpiI3PSa, SpiI3PSb1, SpiI3PSb2 and SpiI3PSc) (Figure S26). The *Lycopersicon* I3PSs protein sequences align well, except for SpiI3PSc, which showed a deletion of 36 amino acid residues (Figure 6a, b). To evaluate if the four *S. pimpinellifolium* proteins are catalytically functional, we used computational 3D molecular structure modeling.

The 3D structures of the four *S. pimpinellifolium* I3PS proteins were inferred with high confidence by homology modeling based on the ~55% identical yeast MIP 1-L-*myo*-inositol-1-phosphate synthase (Stein and Geiger, 2002). When the *S. pimpinellifolium* I3PS sequences were superimposed onto this tetrameric and catalytically competent model structure (1jki) in complex with nicotinamide adenine dinucleotide (NAD), ammonium (NH_4_^+^) and the inhibitor 2-deoxy-glucitol-6-phosphate (DG6), we observed that the short deletions/insertions of one to three residues in SpiI3PSa and SpiI3PSb are distant from the active site (Figures 6a), and thus unlikely to affect the catalytic function. All MIP residues that form the ligand and cofactor binding sites are strictly conserved, except for F307 and G179 (numbering based on SpiI3PSa), which replace, respectively, a leucine and a serine in MIP (Figures 6a and c). Analysis of the homology models strongly suggested that these two substitutions can be accommodated by the 3D environment and do not affect binding and turnover of NAD (Figure 6c). Conversely, SpiI3PSc showed a deletion of 36 residues (red regions in Figure 6b) that could potentially affect the structure of the “lid” that covers the site that binds NAD. While this deletion might not completely abolish catalytic activity, it may result in a lower affinity for NAD and/or a more rapid (but possibly less efficient) substrate turnover. The dimerization and tetramerization interface of yeast MIP was intact and preserved in Spil3PSc, suggesting that all of the I3PS enzymes in *S. pimpinellifolium* form stable and functional tetramers. We therefore conclude that the I3PS enzymes in *S. pimpinellifolium* are functional; thus, supporting the critical relevance of I3PS catalytic function.

**Figure 6.**
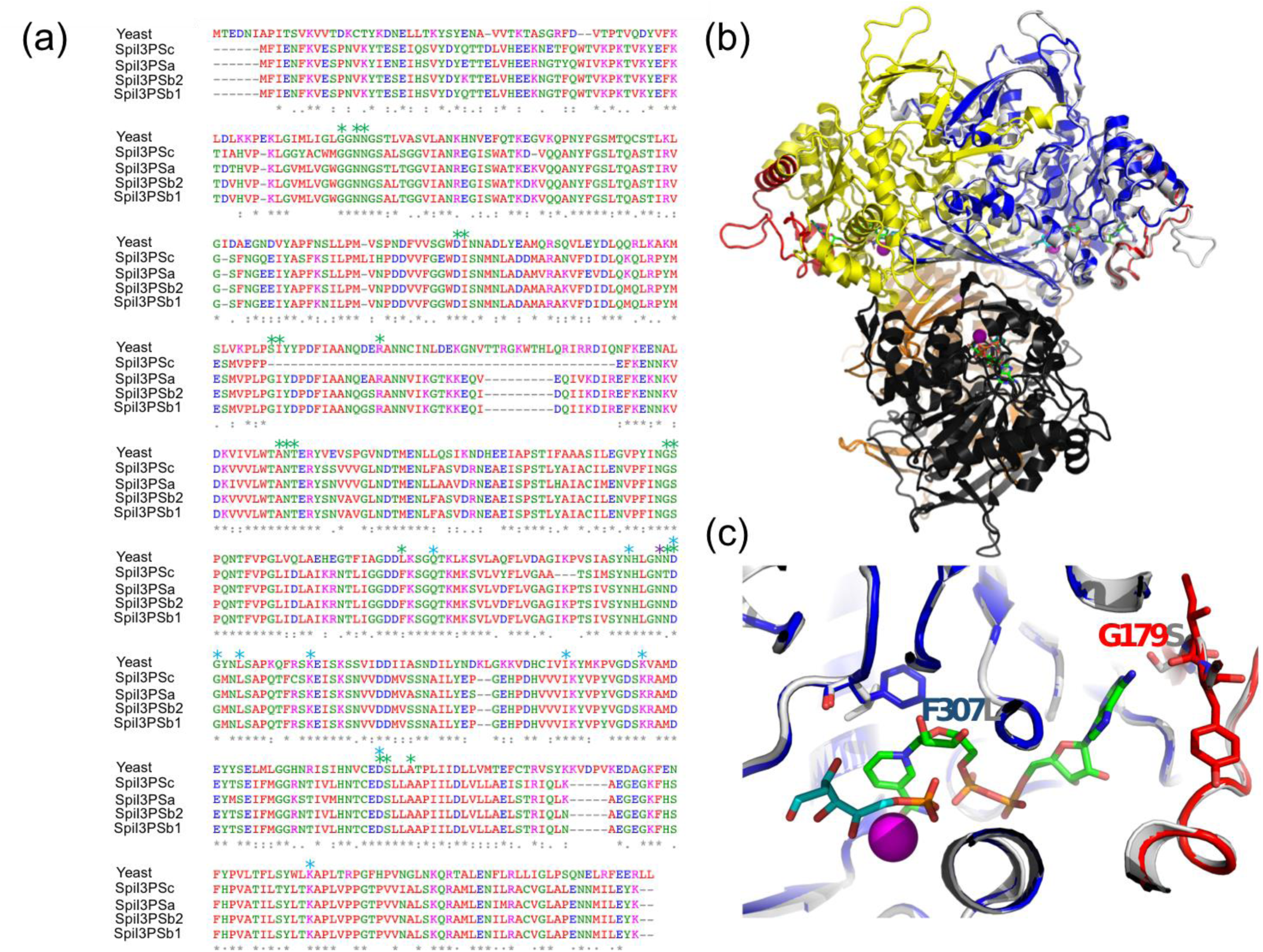
Structural evaluation of the catalytic activity of *S. pimpinellifolium* I3PS proteins. A) Multiple sequence alignment. Yeast MIP protein. Asterisks label residues involved in binding to NAD (green), DG6 (cyan) and NH_4_ (magenta). B) Overall view of the MIP tetramer (PDB: 1jki); individual monomers are shown in gray and yellow (dimer A) and orange and black (dimer B). Red: regions deleted in SpiI3PSc. Blue: homology model of SpiI3PSa superposed. NAD is shown as stick model with green carbons, and DG6 as stick model with cyan carbons, and NH_4_ as magenta sphere; C) Detail of the binding site. Colors as in (B). Side chains discussed in the text are shown.

Given the apparent increase of *I3PS* gene copy number in *S. pimpinellifolium*, we examined the broader inositol-related pathway in *S. pimpinellifolium* using DEAP. We observed that the *S. pimpinellifolium* genome contains all the genes necessary for 1-phosphatidyl-1D-*myo*-inositol and *myo*-inositol cycling according to the inositol phosphate metabolism reference pathway in KEGG (http://www.genome.jp/kegg/pathway/map/map00562.html, verified on 20^th^ of July, 2017). The two key enzymes involved in these cycling processes are Inositol 3-kinase (EC:2.7.1.64) and CDP-diacylglycerol-inositol 3-phosphatidyltransferase (EC:2.7.8.11) (Figure 7), with both enzymes regulating the speed at which the central compound, *myo*-Inositol, and its derivatives are produced. We also observed that the two entry points into the inositol pathway are present in *S. pimpinellifolium* inositol pathway, namely: 1) I3PS (EC:5.5.1.4), which catalyzes the conversion of Glc6P to D-*myo*-inositol-3-phosphate, and 2) phosphatidylinositol-3-phosphatase enzyme (EC:3.1.3.64), which catalyzes the conversion D-*myo*-inositol 1,3-bisphosphate to *myo*-inositol 1-phosphate. Thus, the inositol phosphate metabolism pathway in *S. pimpinellifolium* appears to be complete in terms of entry points and main compounds required for the synthesis of *myo*-inositol. The gene copy-number between species is the same for the majority of the enzymes present in the inositol pathway, with the notable exceptions being I3PS (EC:5.5.1.4), inositol-phosphate phosphatase (EC: 3.1.3.25) and phosphatidylinositol 4-kinase (EC: 2.7.1.67), which have higher gene copy-number in *S. pimpinellifolium* relative to *S. lycopersicum* and *S. pennellii* (Figure 7). These changes may not only lead to an increased *myo*-inositol content but also to modulation of the pattern and concentration of phosphatidylinositols and soluble polyphosphoinositols. Moreover, changes in inositol metabolism will likely impact the concentration of other metabolites (Liu *et al.*, 2013; Kusuda *et al.*, 2015).

To assess the expression level of the inositol metabolism pathway genes we explored RNA-seq on leaf samples from *S. pimpinellifolium* plants grown under control or saline conditions (Table S19). To normalize the expression levels of the target genes, we examined several tomato reference genes (Table S20), and selected tubulin-beta as an adequate reference gene on the basis of its stability between treatments (Figure S28). Our analysis suggested that *I3PS* (EC:5.5.1.4) and inositol-1,4,5-trisphosphate 5-phosphatase gene (EC:3.1.3.56) are up-regulated under salt stress in *S. pimpinellifolium* (Tables S21 and S22). In cultivated tomato, a previous study showed that *myo*-inositol production increases under salt stress (Sacher and Staples, 1985); however, to our knowledge, there is no available expression data of the key genes involved in the inositol pathway under salt stress in this species. The up-regulation of *I3PS* under salinity has been observed in previous studies using *Lotus japonicus* (Sanchez *et al.*, 2008a), *Mesembryanthemum crystallinum* (Nelson *et al.*, 1998) and *Populus euphratica* (Brosché *et al.*, 2005). The increased accumulation of inositol under salinity stress has been observed in salt-tolerant species such as *Eutrema salsugineum* (formerly known as *Thellungiella halophila*), relative to the closely related *Arabidopsis thaliana* (Gong *et al.*, 2005). This metabolic response resulted from increased expression levels of genes involved in the inositol pathway (Gong *et al.*, 2005). The presence of higher levels of inositol in salt-tolerant species has been suggested as an adaptive response of salt-tolerant species by a metabolic anticipation of stress (Sanchez *et al.*, 2008b).

Next, we analyzed the *myo*-inositol content in the leaf tissues of ‘LA0480’ and ‘Heinz 1706’ from hydroponically grown plants, under control and saline conditions. Our results showed a significant increase in the amount of *myo*-inositol produced under saline conditions in both species, but no significant difference in this response was observed between *S. pimpinellifolium* and *S. lycopersicum* (Figure S30 – bottom panels). The quantification of *myo*-inositol in both species was unable to shed light on the importance of the extra copy-numbers of *I3PS* in *S. pimpinellifolium*. Thus, we hypothesize that the higher salinity tolerance of *S. pimpinellifolium* may be related to differences in expression or function of downstream compounds in the inositol pathway, such as different polyphosphoinositides that are involved in signaling pathways, or differences in D-glucuronate that leads to sugar interconversions and/or ascorbic acid levels. For example, in Arabidopsis, the overexpression of the purple acid phosphatase gene (*AtPAP15*), a phytase that hydrolyzes phytate to *myo*-inositol and free phosphate, led to the accumulation of ascorbic acid in the shoot and an increase in salinity tolerance (Zhang *et al.*, 2008). Additionally, ^downstream inositol derivatives such as Ins(1,4,5)*P*_3_, PtsIns(4,5)*P*_*2*_ and PtdIns*4P* (Figure 7) have been^ shown to play a role in abiotic stress signaling (reviewed by Munnik and Nielsen, 2011). For instance, Ins(1,4,5)*P*_3_, has been suggested to contribute to drought tolerance in tomato (Khodakovskaya *et al.*, 2010), and could also play a role in salinity tolerance. Furthermore, *phosphoinositide phospholipase C* (EC:3.1.4.11) expression has been shown to increase in response to salinity stress in both rice and Arabidopsis and is required for stress-induced Ca^2+^ signaling and for controlling Na^+^ accumulation in leaves (Munnik and Nielsen, 2011; Li *et al.*, 2017). Similarly, in *S. pimpinellifolium*, the higher copy number of *phosphatidylinositol 4-kinase* (EC:2.7.1.67) (Figure 7 and Table S18) and the increased expression of *1-phosphatidylinositol-4-phosphate 5-kinase* (EC:2.7.1.68), *phosphatidylinositol phospholipase C* (EC:3.1.4.11) as well as *phosphatidylinositol 4-kinase* (EC:2.7.1.67) (Table S22) may be involved in the increased salinity tolerance of *S. pimpinellifolium*.

Because the accumulation of *myo*-inositol in the cytoplasm of cells under stress is thought to be related to the tissue tolerance mechanism (Nelson *et al.*, 1999; Munns, 2005; Roy *et al.*, 2014), we investigated the Na and K concentration in the same tissues (Figure S30, top and middle panels). We observed that Na accumulates to higher levels in *S. pimpinellifolium* compared to *S. lycopersicum*, yet *S. pimpinellifolium* is more salt-tolerant; thus, reinforcing the idea that tissue tolerance is the main mechanism of salinity tolerance in this species. Interestingly, other tomato wild relatives besides *S. pimpinellifolium*, namely *S. pennellii*, *S. peruvianum* and *S. galapagense* also accumulate higher concentrations of Na while being more salt-tolerant than cultivated tomato (Tal, 1971; Santa-Cruz *et al.*, 1999; Almeida *et al.*, 2014), which may also suggest that tissue tolerance could be the main mechanism of salinity tolerance in these wild species.

Although much research has been conducted into the biochemistry of inositol-related pathways, we are still far from fully understanding their underlying complexity. Specifically, to our knowledge, the link between these pathway derivatives and stress-response mechanisms have not been fully elucidated. As such, further studies on the role of these derivatives in processes such as Ca^2+^ signaling, osmoprotection and maintenance of membrane integrity are expected to reveal the basis of the higher salinity tolerance of *S. pimpinellifolium* compared with *S. lycopersicum*.

**Figure 7.**
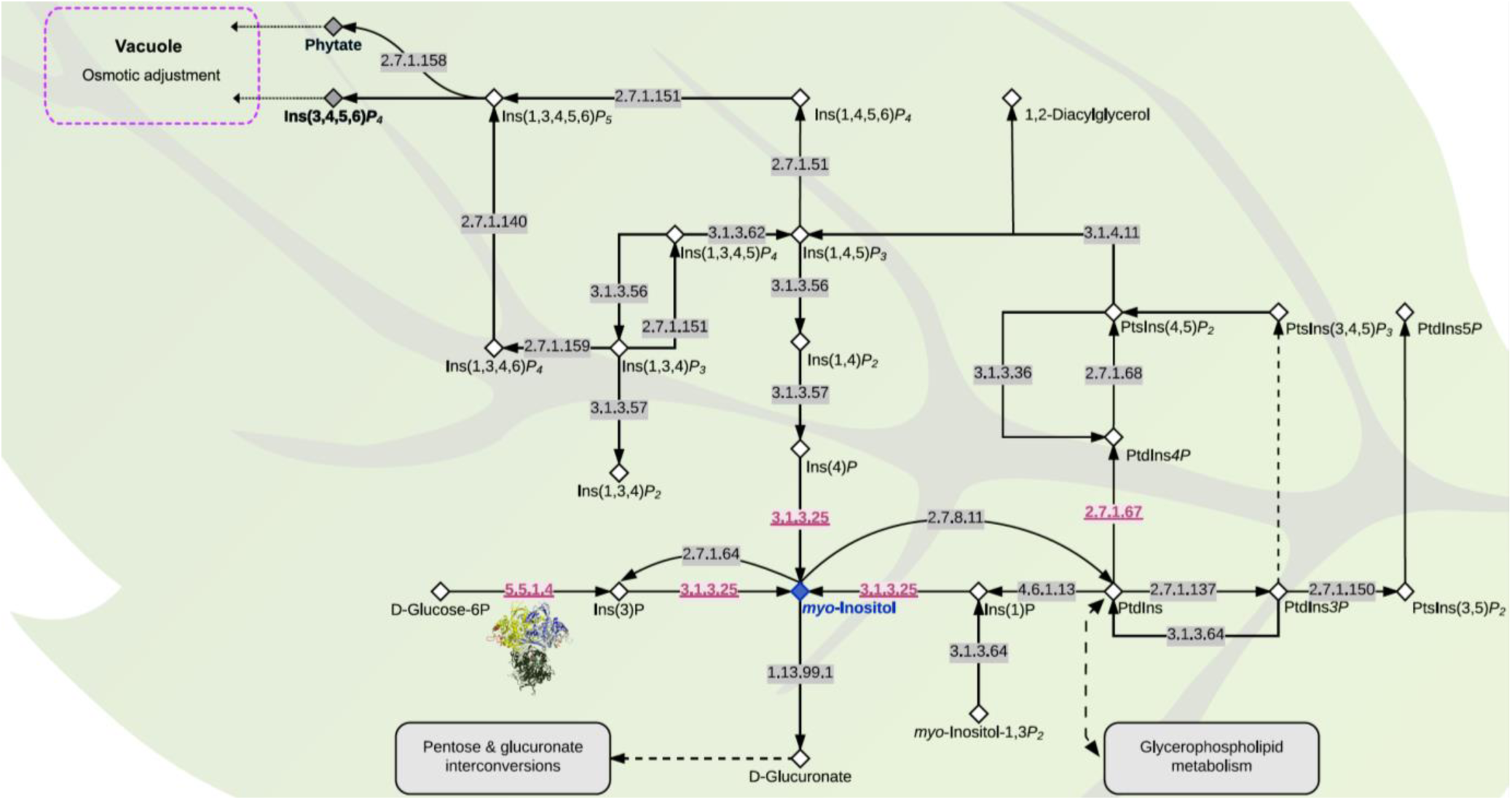
The inositol metabolism pathway in *S. pimpinellifolium* and *S. lycopersicum*. The pathway was adapted from the KEGG inositol metabolism pathway (map00562-http://www.genome.jp/kegg/pathway/map/map00562.html). Compounds are represented with diamonds, *myo*-inositol is shown in a blue diamond whereas phytate and Ins(1,3,4,5)*P*_4_ are represented by a grey diamond. Enzymes are represented with their EC numbers placed directly on arrows. Enzymes with gene copy numbers higher in *S. pimpinellifolium* than in *S. lycopersicum* are underlined and colored in red (Table S18). Compound abbreviations were taken from the ChEBI database (Hastings *et al.*, 2013): Ins(1)P: Inositol 1-phosphate; PtdIns: Phosphatidyl-1D-*myo*-inositol; PtdIns3*P*: 1-Phosphatidyl-1D-*myo*-Inositol-3*P*; PtsIns(3,5)*P*_2_: 1-phosphatidyl-1D-*myo*-inositol 3,5-bisphosphate; PtdIns5*P*: 1-phosphatidyl-1D-*myo*-inositol 5-phosphate; PtsIns(3,4,5)*P*_3_: 1-phosphatidyl-1D-*myo*-inositol 3,4,5-trisphosphate; PtsIns(4,5)*P*_2_: 1-phosphatidyl-1D-*myo*-inositol 4,5-bisphosphate; Ins(1,4,5)*P3*: 1D-*myo*-inositol 1,4,5-trisphosphate; Ins(1,3,4,5)*P*_4_: 1D-*myo*-inositol 1,3,4,5-*P*_4_; Ins(1,4,5,6)*P4*: 1D-*myo*-inositol 1,3,4,5-*P*_4_; Ins(1,3,4,5,6)*P5*: 1D-*myo*-inositol 1,3,4,5,6-*P*_5_.

## CONCLUSIONS

*S. pimpinellifolium* has the potential to increase the genetic diversity of cultivated tomato. Despite the availability of a draft genome sequence of *S. pimpinellifolium*, limited progress has been made towards unlocking the genetic potential of this species. Our work provides the basis to accelerate the improvement of cultivated tomato by presenting the genome sequence and annotation of the salt-tolerant *S. pimpinellifolium* accession ‘LA0480’. Our genome analysis shows that *S. pimpinellifolium* is enriched in genes involved in biotic and abiotic stress responses in comparison to cultivated tomato. Moreover, we demonstrate the increased salinity tolerance of ‘LA0480’, and suggest that it could be related to the inositol-related pathways. The expansion of inositol-3-phosphate synthase gene copies in *S. pimpinellifolium*, which encodes a key enzyme in the inositol pathway, may contribute to its higher salinity tolerance when compared to *S. lycopersicum.* Future studies are necessary to validate the role of I3PS in salinity tolerance in tomato, for instance by using genetic tools (e.g. gene knockout and overexpression) and metabolic profiling by quantifying inositol derivatives. Altogether, our work will enable geneticists and breeders to further explore genes that underlie agronomic traits as well as stress-tolerance mechanisms in *S. pimpinellifolium*, and to use this knowledge to improve cultivated tomato.

## EXPERIMENTAL AND BIOINFORMATIC PROCEDURES

### Field trial for assessment of salinity tolerance

The field data presented is a subset from a larger field trial conducted at the International Center for Biosaline Agriculture (ICBA) in Dubai, United Arab Emirates (N 25˚ 05.847; E 055˚ 23.464), between October 2015 and May 2016. We used a randomized block design, with a non-saline and a saline plot, each comprising four blocks, each with at least one randomly positioned replicate from every accession. Plants were planted in rows, with 0.5 m spacing between plants, and 1 m spacing between rows. Plants were grown in a nursery for 6 weeks before transplanting in the field. Both plots were irrigated with non-saline water for the first 5 weeks following transplantation. After 5 weeks, the irrigation for the saline field was switched to a saline source. Regular water sample collection and analysis over the course of the experiment indicated an average electroconductivity (EC) of 0.7 dS/m^2^ and 12.3 dS/m^2^, and a NaCl concentration of 0.5 - 10 mM and 70 - 110 mM for the non-saline and saline water sources, respectively. After salt-stress application, the experiment was continued for 17 weeks. Mature fruit were harvested continually throughout the field trial to assess fruit-and yield-related traits and a final destructive harvest was performed to evaluate biomass traits (Supplementary material). All measurements were spatially corrected in the R statistical computing environment (v2.12), using custom scripts and the ASReml v3.0-1 (Gilmour *et al.*, 2009) package for R v3.2.0 (R Core Team, 2014).

We calculated the Harvest Index (HI) using the formula:

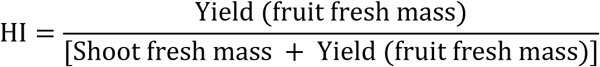

In tomatoes, the HI is the fresh fruit yield as a proportion of the total fresh shoot mass (including fruit), (Gianfagna *et al.*, 1997). Salinity tolerance (ST) was calculated for each trait in each genotype using the formula:

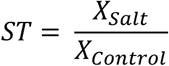

where *X*_*salt*_ and *X*_*control*_ are the mean value of variable *X* under salt stress and control conditions, respectively. ST gives a measure of the plant’s ability to maintain the variable X under salt stress relative to control conditions; explicitly, it provides the value under stress as a proportion of the control value. A ST of less than one suggests that performance under stress is lower than under control conditions while a ST of greater than one suggests that performance under stress is higher than under control conditions

### ‘LA0480’ DNA library construction, sequencing and assembly

The salt tolerant *S. pimpinellifolium* accession ‘LA0480’ was sequenced using the HiSeq 2000 Illumina platform at King Abdullah University of Science and Technology (KAUST) (Figure 8). DNA was extracted from whole flowers collected from a single soil-grown plant ‘LA0480-ref’ using the Qiagen DNeasy Plant Mini Kit (Qiagen, Germany), following manufacturer’s guidelines. Two paired-end short-read libraries (139 and 332 bp mean insert length) and five mate-pair libraries (2, 6, 8, 10 and > 10 kb target insert length) were prepared using the NEBNext Ultra DNA Library Prep Kit and the Nextera Mate-pair Library Kit, respectively (New England Biolabs, UK).

**Figure 8.**
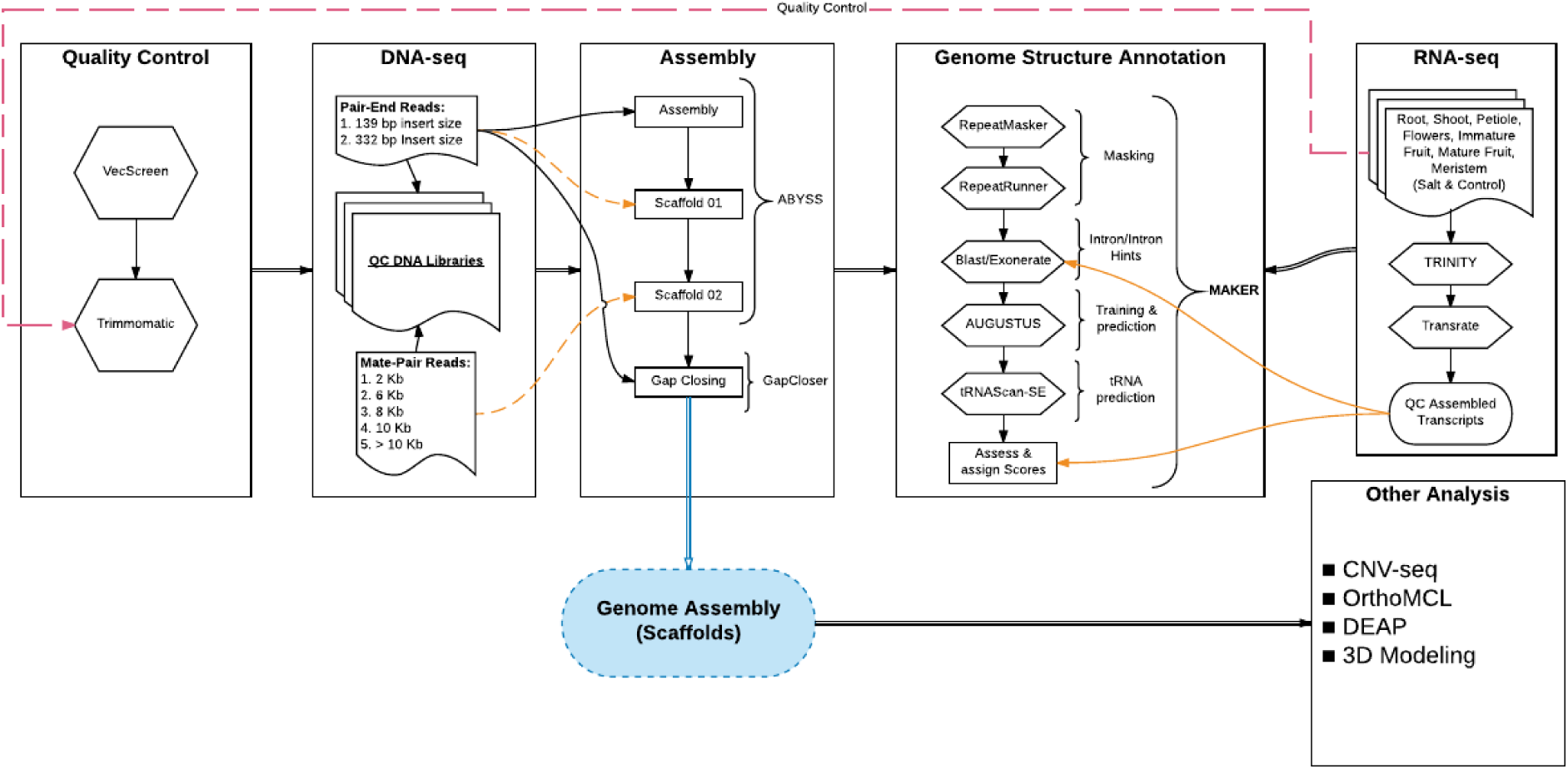
Schematic overview of the main tools used for the genome sequencing and assembly of *S. pimpinellifolium* ‘LA0480’. The diagram outlines the workflow and the main tools that were used in the different stages of assembly, gene model annotation and functional annotation.

Adapter sequences, low-quality four nucleotide stretches of nucleotides, and low quality leading and trailing bases were removed with Trimmomatic v0.33 (Bolger *et al.*, 2014b) and reads with final a final length of less than 36 bp after trimming were discarded (Table S1). Quality of the data was verified before and after trimming with FastQC (Andrews, 2010).

Processed pair-end data were *de novo* assembled into contigs using ABySS (Simpson *et al.*, 2009). Default parameters were used apart from a k-mer length of 77 bp, which was determined based on a kmer analysis (Figure S1). Initial contigs were scaffolded based on library size information of the PE reads libraries (Table S2), followed by a second round of scaffolding with mate pair data utilizing the ABySS pipeline. Preliminary quality control was performed by mapping the sequencing reads back to the genome with bwa (bwa mem) (Table S3). GapCloser (Luo *et al.*, 2012) was used to close gaps in the assembled scaffolds (Table S4). The completeness of the genome assembly was assessed with BUSCO (Simão *et al.*, 2015). Assembly quality was also assessed by mapping the RNA-seq data onto the assembled scaffolds. The GC content was estimated based on the raw data using FastQC and based on the assembled scaffolds using fasta_tool (https://github.com/The-Sequence-Ontology/GAL).

### ‘LA0480’ transcriptome sequencing and assembly

RNA was extracted from root, young leaf, old leaf, petiole, meristem, flower, and immature fruit tissue samples collected from the mature soil-grown ‘LA0480-ref’ plant. Additionally, leaf and root samples from plants (‘LA0480-ref’ progeny) grown hydroponically under control and salt stress conditions were collected (Tables S6 and S7). RNA was extracted using ZR Plant RNA MiniPrep Kit (Zymo, CA, USA). RNA sequencing libraries were prepared using the NEBNext Ultra Directional RNA Library Prep Kit for Illumina (New England BioLabs, UK). Sequencing reads were processed with Trimmomatic, and assembled into transcripts using Trinity v2.0.6 (Grabherr *et al.*, 2011).

### Analysis of repetitive elements

RepeatModeler v1.0.8 and RepeatMasker v4.0.5 (Tarailo-Graovac and Chen, 2009) were used to screen the *S. pimpinellifolium* genome for repetitive elements (RE). A library of *de novo* repeats was constructed with RepeatModeler and this library was subsequently merged with the RepBase library (v21.02) from RepeatMasker. RepeatMasker was run on the assembled genome (minimum length of 5 kb) using the total repeat library.

### ‘LA0480’ gene structure annotation

To call genes and their structure, we used the MAKER annotation pipeline v03 (Cantarel *et al.*, 2008) with AUGUSTUS (Stanke *et al.*, 2004) as the base *ab initio* gene predictor. AUGUSTUS was trained using the existing *S. lycopersicum* gene model as the basis and the assembled RNA-seq data as hints (see supplemental material for more on AUGUSTUS training). Before gene calling, repeat elements were masked using RepeatMasker and RepeatRunner. Protein-coding genes were predicted using hints from the assembled transcripts as well as from the unassembled raw RNA-seq data and from the aligned proteins from *S. lycopersicum*, *S. pennellii* and SwissProt (Bairoch and Apweiler, 2000). tRNA genes were predicted using tRNAscan-SE (Lowe and Eddy, 1997). The predicted genes were assessed and assigned scores using MAKER based on the assembled transcripts and homologous proteins. For a detailed description of the annotation, refer to the supplementary material.

Functional annotation was performed using DEAP. KEGG Orthologs (KO) were assigned based on the KEGG database using BLASTp with the following parameters (BLAST percent identity cut off of 60, maximum *E*-value of 1E-5). Functional domains, protein signatures and their associated Gene Ontology (GO) were assigned using InterProScan (Jones *et al.*, 2014). For versions of the different tools and databases used under DEAP v1.0 refer to http://www.cbrc.kaust.edu.sa/deap/. For a detailed description of the annotation, refer to the supplementary material.

### Identification of orthologous genes

Orthologous and paralogous protein relationships between the four species were identified using OrthoMCL (Li *et al.*, 2003). All-against-all BLASTP comparisons (Blast+ v2.3.0) (Camacho *et al.*, 2009) were performed using recommended settings. Custom Perl scripts were utilized to analyze OrthoMCL outputs for visualization with InteractiVenn (Heberle *et al.*, 2015). Protein datasets for *S. pennellii* and *S. lycopersicum* were obtained from the Sol Genomics Network (https://solgenomics.net/). The *S. tuberosum* protein dataset was obtained from Phytozome (https://phytozome.jgi.doe.gov). All sequences were downloaded in February 2017. The proteins corresponding to the primary transcripts were identified with custom Perl scripts.

### CNV-seq and SNP analyses

CNV were investigated using CNV-seq v0.2.7 (Xie and Tammi, 2009). Raw reads from *S. pimpinellifolium* and *S. lycopersicum* (SRR404081) were aligned to the *S. lycopersicum* reference genome (NCBI assembly accession GCF_000188115.3) using BWA v0.7.10 (Li and Durbin, 2009). Alignment files were post-processed using SAMtools v1.3.1 (Li *et al.*, 2009). Then, short reads data from *S pimpinellifolium* and *S. lycopersicum* were mapped to the *S. lycopersicum* genome using the following settings: p ≤ 0.001, log_2_ threshold ≥ +/-1, window size= 276, minimum window of 4 and using the *S. lycopersicum* genome-size of 813 Mb. The circular plot was generated using CIRCOS v0.69.3 (Krzywinski *et al.*, 2009). We also produced high and low CNVs graphs for all 12 chromosomes using R (R Core Team, 2014) (Figure S7).

For SNPs analysis, the short read sequence data from *S. pimpinellifolium* were mapped to the *S. lycopersicum* reference genome as described in the CNV section above. SNPs were called using the mpileup command of SAMtools (v1.3.1) and custom Perl scripts were used to filter SNPs for a depth of at least 8 and a SNP allele frequency greater than 75%. SNPs were binned into 1 Mb bins, and plotted together with the CNV data using CIRCOS.

### KO enrichment analysis

KO enrichment analysis for the *S. pimpinellifolium* and *S. lycopersicum* genomes was performed using DEAP Compare. For a detailed description of the comparison, refer to the supplementary material. Only KO terms that were assigned based on BLAST percentage identity of at least 60% and above were considered (*E*-value ≤ 1E-5). For each observed KO_i_, we compared the ratio KO_i_ / KO_total-observed_ in each species using Fisher’s exact test (confidence interval 0.95). An enrichment is defined where the *P* value is significant (*P* < 0.05). We corrected for multiple testing using the Benjamini-Hochberg method.

### Identification of salt tolerance candidate genes and orthologs in tomato species

The salt tolerance candidate gene list was adapted from Roy *et al.* (2014) (Tables 3 and S15) and verified against 'Dragon Explorer of Osmoprotection associated Pathways' - DEOP (Bougouffa *et al.*, 2014). For CGs with supporting literature in *S. pimpinellifolium*, protein sequences were compared using BLASTp and multiple sequence alignment (MSA) tools such as MUSCLE (Edgar, 2004) or KAlign (Lassmann and Sonnhammer, 2005). We also performed BLASTp searches (setting a high percentage identity, i.e. thresholds usually > 90%) and used OrthoMCL orthogroups to verify the orthology. For CGs with no supporting literature in *S. pimpinellifolium*, we investigated CG orthologs in *S. lycopersicum* using a combination of approaches: 1) BLASTp against *S. lycopersicum* total proteins; 2) orthogroup identification using OrthoDB; 3) inspection and comparison of functional domains; and 4) MSA and visual assessment of the alignment. Alignments for the CGs are presented in Figures S10-24. The workflow is summarized in the Figure S9. For more details on the methods please refer to Supplementary Material.

### Phylogenetic analysis

The online tool Phylogeny.fr (Dereeper *et al., 2008*) was used for the phylogenetic analysis of seven Solanaceae species, namely the three *Lycopersicon* species *S. pimpinellifolium*, *S. lycopersicum* and *S. pennellii*, potato (*S. tuberosum*), eggplant (*S. melongena*), hot pepper (*Capsicum annuum*) and tobacco (*Nicotiana tabacum*), with *A. thaliana* set as an outgroup. Multiple sequence alignment of the DNA sequences of I3PS from these species was performed using ClustalOmega with two combined guide-trees and HMM iterations (Sievers *et al.*, 2011). Details for DNA sequences can be found in Table S18. The construction of the phylogenetic tree was estimated using the maximum likelihood method (PhyML), and the Generalized Time Reversible substitution model (GTR) (bootstrap value = 100). The tree was drawn with TreeDyn (Chevenet *et al.*, 2006).

### Computational structural analysis of I3PS proteins

SwissModel (Arnold *et al.*, 2006) was used to produce homology models based on the ~55% identical structure of the yeast MIP 1-L-*myo*-inositol-1-phosphate synthase [PDB id 1jki (Stein and Geiger, 2002); QMEAN values are between -2.0 and -2.3 for SpiI3PSa and SpiI3PSb alleles, and -3.2 for SpiI3PSc]. Models were manually inspected, and mutations evaluated, using the Pymol program (pymol.org).

### *Myo*-inositol content determination

The selected tissues, old leaf (youngest fully expanded leaf at the time of salt imposition) and young leaf (youngest fully expanded leaf at the time of harvest) were harvested from seedlings grown following the “Hydroponics 2” protocol (Supplementary material) seven days after salt stress application. Frozen leaf samples were ground, freeze-dried and 20 mg of tissue was mixed with water. After centrifugation, *myo*-inositol content in supernatant was measured using K-INOSL assay kit according to the manufacturer's instructions (Megazyme International Ireland, Bray, Wicklow, Ireland).

### Measurement of shoot ion concentration

Young and old leaves were collected from plants grown in parallel to those prepared for *myo*-inositol quantitation to assess the concentration of Na and K in leaf tissues. The fresh and dry mass of each sample was measured to determine the tissue water content. Dried leaf samples were digested overnight in 1% (v/v) nitric acid (HNO_3_) in an oven at 70°C. The concentrations of Na and K were determined in three biological replicates using a flame photometer (model 420; Sherwood Scientific Ltd., Cambridge, UK).

## ACKNOWLEDGEMENTS

This publication is based upon work supported by the King Abdullah University of Science and Technology (KAUST) Office of Sponsored Research (OSR) under Award No. 2302, No. 1976-02 and KAUST Base Research Funds to VBB grant No. BAS/1/1606-01-01. Genome sequencing was performed at the biological core laboratories of KAUST. All the computational analyses were performed on Dragon and Snapdragon computer clusters of the Computational Bioscience Research Center (CBRC) at King Abdullah University of Science and Technology (KAUST). We thank Gabriele Fiene (KAUST) for her assistance with the field trial and phenotypic data collection. We thank Hajime Ohyanagi for his comments on the phylogenetic analysis.

## ACCESSION NUMBERS

All raw data for the DNA-seq and RNA-seq are available upon request.

